# Optimising CAR-T cell sensitivity by engineering matched extracellular sizes between CAR/antigen and CD2/CD58 adhesion complexes

**DOI:** 10.1101/2025.01.06.631424

**Authors:** Jake Burton, Jesús A. Siller-Farfán, Violaine Andre, Edward Jenkins, Michael I. Barton, Sofia Bustamante Eguiguren, Jose Cabezas Caballero, Simon J. Davis, Thomas R Weikl, P. Anton van der Merwe, Omer Dushek

## Abstract

Chimeric antigen receptor (CAR)-T cells exhibit low antigen sensitivity, which restricts their therapeutic efficacy and leads to patient relapses when cancer cells downregulate antigen expression. Despite the pressing need to overcome this limitation, the underlying mechanisms remain poorly understood. Here, we demonstrate that enhancing CAR sensitivity to match the sensitivity of the T-cell receptor (TCR) can be achieved by engineering matched extracellular sizes of CAR/antigen and CD2/CD58 complexes. We find that different CAR/antigen sizes, which are generated by different CAR architectures and different target antigens, require a different CD2/CD58 size to optimise sensitivity. This extracellular size-matching improves antigen engagement and co-localisation of CAR/antigen and CD2/CD58 complexes. We also find that size-matching controls co-inhibition of CARs by PD-1/PD-L1. These findngs highlight the importance of size-matching for signal integration by surface receptors and offers a new approach to tune CAR-T cell sensitivity by matching or mismatching extracellular sizes.

**One sentence summary:** The antigen sensitivity of CAR-T cells can be tuned to match the sensitivity of TCR-T cells by varying the relative extracellular size of CAR/antigen and CD2/CD58 complexes.

**Graphical abstract:** 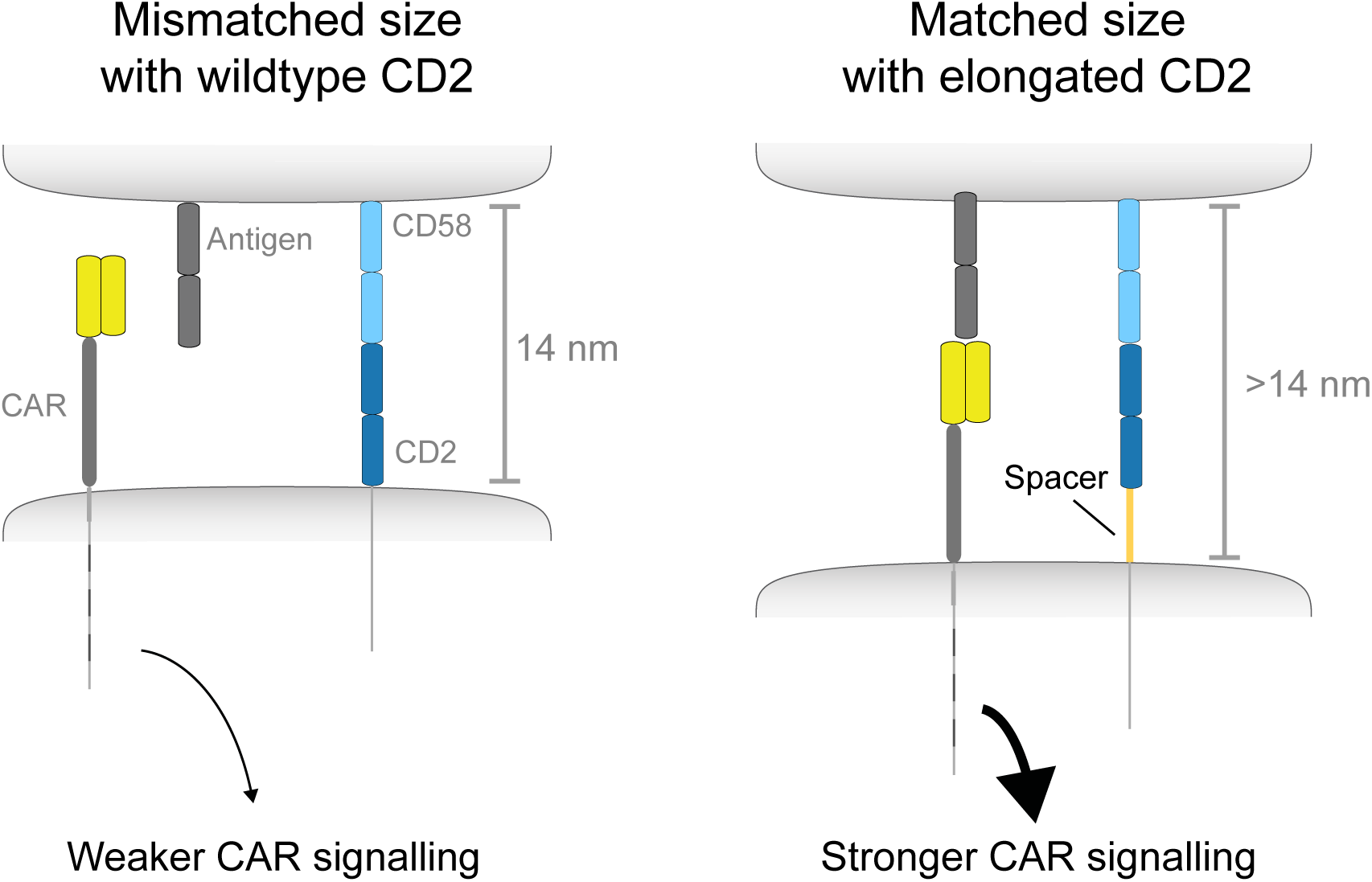

## Introduction

Chimeric antigen receptor (CAR)-T cells targeting CD19 have revolutionised the treatment of B cell malignancies, but their success in other cancers has been limited (1). One reason for this is a major defect in their antigen sensitivity; CARs require *>*100-fold more antigen to activate T cells than the T cell receptor (TCR) (2–5). This low sensitivity has hampered the wider deployment of CAR-T cells to tumours expressing low levels of antigen, and it contributes to patient relapse when tumour cells re-emerge with lower levels of antigen (6–9).

The mechanism underlying this antigen sensitivity defect in CARs remains obscure. Several studies have shown enhanced sensitivity by chimeric receptors that integrate the antigen binding domains of CARs into the TCR-CD3 complex (e.g. STARs, HITs, and TRuCs (4, 10–13)). This suggests that high sensitivity relies on the full complement of TCR-CD3 signalling, which is lacking in standard clinically approved CARs. However, as these chimeric receptors also alter extracellular structure relative to standard CARs, it is unclear whether intracellular signalling or extracellular structure is responsible for their enhanced sensitivity. Indeed, studies have shown that changing the extracellular structure of standard CARs can impact their sensitivity (14–17). For example, the higher sensitivity of 2nd generation CARs that use CD28 rather then 4-1BB signalling has been traced to their respective CD28 or CD8a extracellular hinge (15). Therefore, it remains unclear whether extracellular structure or intracellular signalling determines antigen sensitivity. Moreover, improvements in CAR sensitivity have remained below the very high antigen sensitivity of the TCR (18–20).

T cells achieve high antigen sensitivity through their TCRs partly by exploiting other surface receptor/ligand interactions (20). The adhesion receptor CD2 binding to its ligand CD58 has been shown to be important for TCR recognition of pMHC in the context of infection and cancer (21–25). In reductionist systems, CD2 engagement can dramatically increase the sensitivity of the TCR by ∼ 100-200-fold but has only a modest ∼ 5-10-fold impact on CAR sensitivity (4, 5). It remains unclear why CARs fail to efficiently exploit CD2 adhesion.

Here, we study the impact of changes in the extracellular size of CAR/antigen and CD2/CD58 complexes on antigen sensitivity. We find that engineering matched sizes of these complexes can fully restore the antigen sensitivity of CARs to match those of the TCR, and that this is accompanied by increased antigen engagement and closer co-localisation between CAR/antigen and CD2/CD58 complexes within the immune synapse. We hypothesise that size-matching underlies signal integration by other co-signalling receptor/ligand interactions and show that increasing the size of the inhibitory receptor PD-1 reduces its activity on the TCR but increases its activity on CARs. Collectively, our work highlights that CARs do not require integration into the TCR-CD3 complex to achieve high sensitivity and offers a new size-matching approach to engineering desired levels of CAR-T cell sensitivity.

## Results

### A compact CAR efficiently exploits CD2/CD58 adhesion and achieves high antigen sensitivity

Chimeric receptors in the STAR (also known as HIT) and the TRuC formats are expected to induce the full complement of TCR-CD3 signalling but they also alter the extracellular structure/size relative to standard CARs (Fig. 1A). To investigate whether extracellular size alone can restore sensitivity, we designed a two-chain Compact CAR that fused the Fab heavy and light chains to the transmembrane and cytoplasmic domain of CD28, followed by the *ζ*-chain (Fig. 1A, Fig. S1). This Compact CAR combines the overall extracellular size of the TCR and STAR (2 Ig domains) with the transmembrane and cytoplasmic signalling regions of standard CARs enabling us to identify whether signalling or size determines antigen sensitivity.

**Figure 1:**
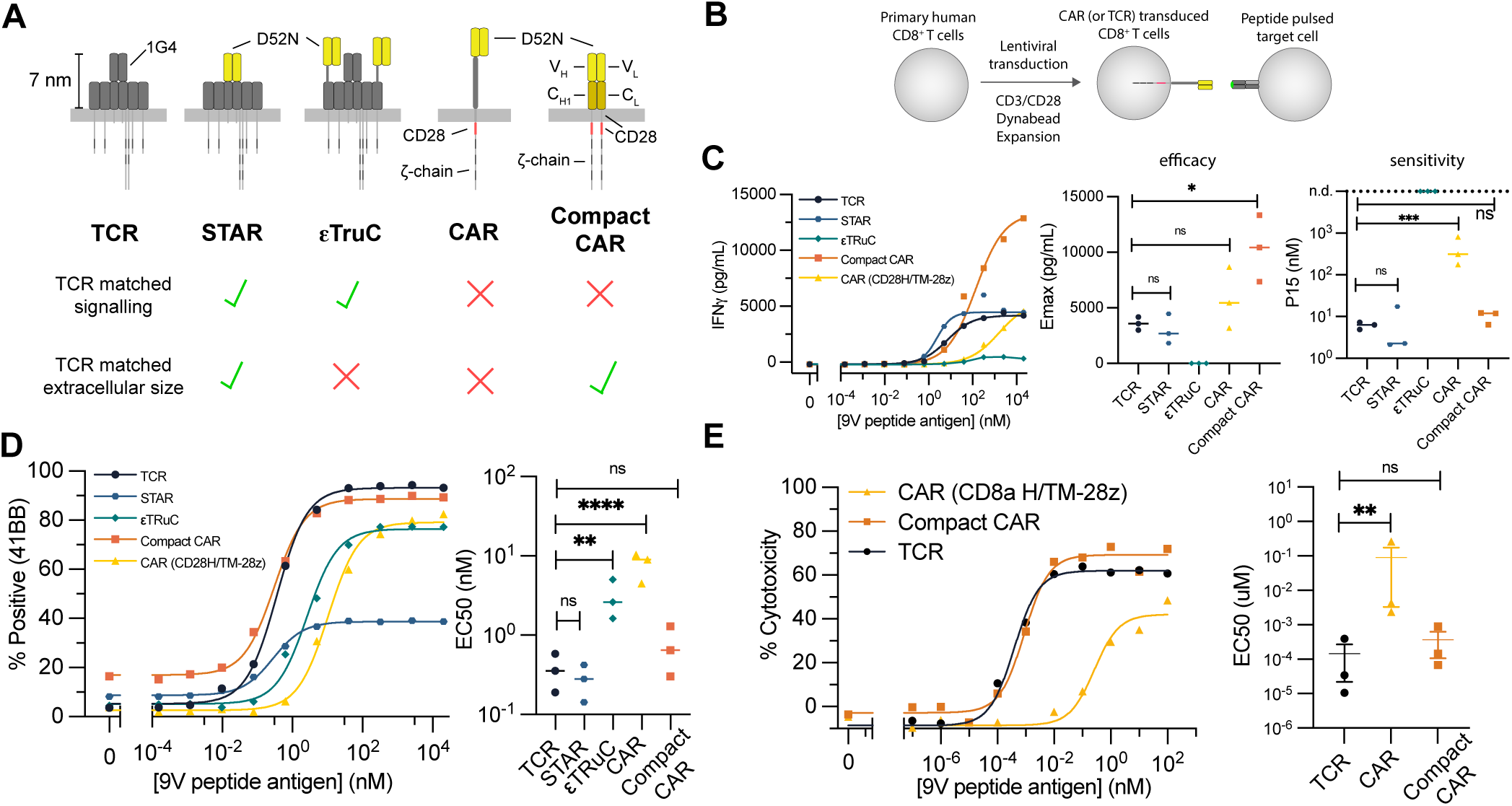
A Compact CAR that is independent of the TCR-CD3 complex achieves the same antigen sensitivity as the TCR. **(A)** A Compact CAR shares extracellular dimension but not signalling with the TCR. The chimeric receptors rely on the D52N scFv to recognise the NY-ESO-1 peptide displayed on HLA-A*02:01 (26). **(B)** Sensitivity is compared by peptide antigen titrations on U87 target cells. **(C)** Representative IFN-g cytokine and summary measures across N=3 independent experiments (see Fig. S3 for IL-2). **(D)** Representative 4-1BB surface activation marker and summary measures across N=3 independent experiments. **(E)** Representative target killing and summary measures across N=3 independent experiment. A t-test with Holm-Sidak multiple comparison correction compares E_max_ (directly) or EC_50_ (log-transformed). Abbreviations: * = p-value≤0.05, ** = p-value≤0.01, *** = p-value≤0.001, **** = p-value≤0.0001.

Expression of the TCR, STAR, TRuC, Compact CAR, and a standard CAR that recognise the NY-ESO-1 pMHC antigen could readily be detected at the surface of primary human CD8^+^ T cells with the Compact CAR displaying matched expression to a standard CAR (Fig. S2A). We next confirmed that the Compact CAR does not rely on the TCR-CD3 complex by showing that, like standard CARs, it can be detected at the cell surface of CD3^−^ Jurkat T cells (Fig. S2B). Remarkably, the antigen sensitivity of the Compact CAR was far higher than a standard CAR and indistinguishable from the TCR, using multiple T cell activation readouts (Fig. 1C-E, Fig. S3). Additionally, the Compact CAR achieved higher efficacy (maximum cytokine levels) compared to the TCR.

We hypothesised that the Compact CAR achieves higher sensitivity than a standard CAR because it can more efficiently exploit the CD2/CD58 adhesion interaction. To test this hypothesis, we used our recently described CombiCell platform to titrate pMHC alone or in combination with CD58 directly on the cell surface (5) (Fig. 2A). We found that the Compact CAR, like the TCR and in contrast to a CAR, efficiently exploits CD2/CD58 interactions to increase its antigen sensitivity (Fig. 2B-C, Fig. S4). Taken together, these results suggest that the inefficient use of CD2/CD58 co-stimulation and the low sensitivity of standard CARs is primarily a result of differences in the CAR and TCR extracellular domains, rather then differences in their signalling components.

**Figure 2:**
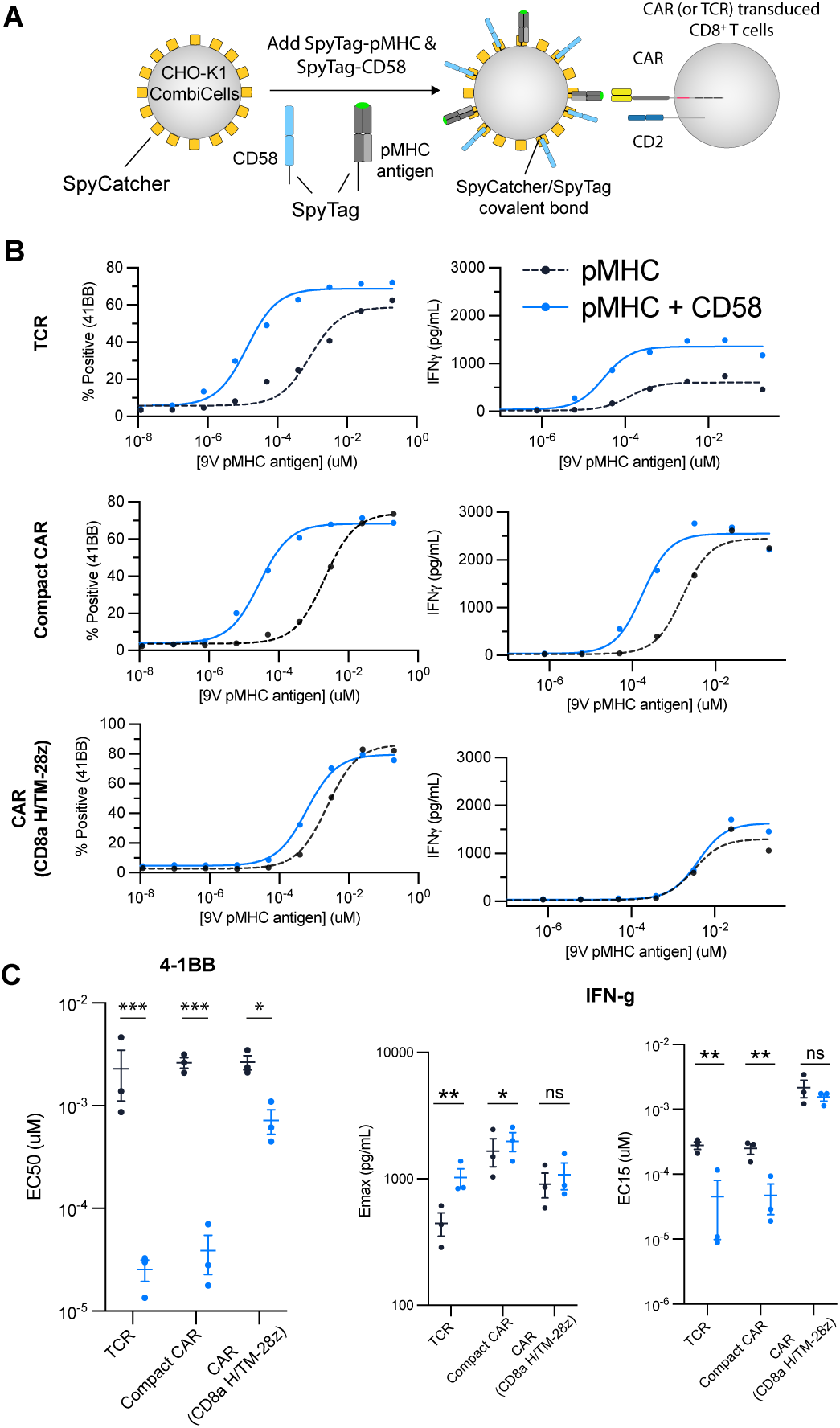
The Compact CAR efficiently exploits CD2/CD58 co-stimulation. **(A)** CHO-K1 CombiCells expressing surface Spycatcher were coupled with a titration of Spytag-pMHC with or without Spytag-CD58 (0.1 *µ*M) before co-culturing with primary human CD8+ T cells (5). **(B)** Representative activation marker (4-1BB, left column) and cytokine (IFNg, right column) along with summary measures (N=3, bottom) for the indicated receptor. See Fig. S4 for additional readouts. A t-test with Holm-Sidak multiple comparison correction compares E_max_ (directly) or EC_50_ (log-transformed). Abbreviations: * = p-value≤0.05, ** = p-value≤0.01, *** = p-value≤0.001, **** = p-value≤0.0001.

### Optimising the antigen sensitivity of different CARs by engineering nanoscale CD2 elongations

The ability of the Compact CAR to exploit CD2/CD58 with the same efficiency as the TCR suggested that the CD2/CD58 dimensions are matched for the Compact CAR and TCR but not for standard CARs when they bind the same pMHC antigen. While there is no structure for the extracellular hinge of standard CARs, our finding that reducing the size of their hinge increases their sensitivity suggests that they form a CAR/antigen complex that is too large when recognising a pMHC antigen (Fig. S5). This hypothesis predicts that elongating CD2 should enhance antigen sensitivity of CARs but reduce it for TCRs (Fig. 3A).

**Figure 3:**
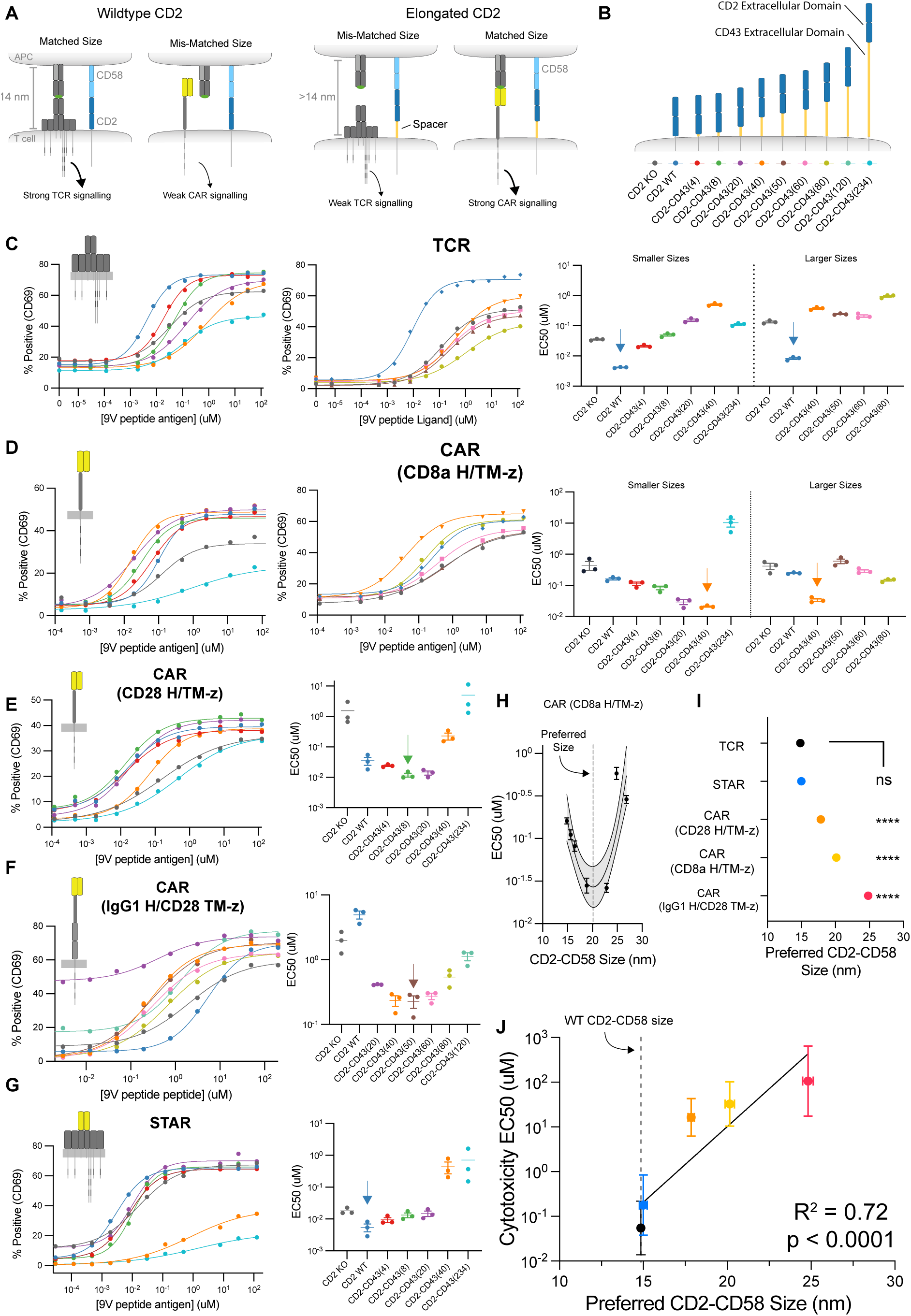
Engineering synthetic elongations of CD2 increases the antigen sensitivity of CAR-T cells. **(A)** Cartoon displaying the hypothesis that matched dimensions between CD2/CD58 adhesion and the antigen receptor/antigen complex determines antigen recepror signalling. **(B)** Cartoon of elongated CD2 molecules. **(C-G)** TCR*α*^−^*β*^−^CD2^−^ E6.1 Jurkat T cells expressing the indicated antigen receptor and CD2 molecule were co-cultured with the U87 cell line for 20 hours. Representative dose-response curves and summary EC_50_ values from N=3 independent experiments. **(H)** The EC_50_ over the estimated CD2-CD58 size is fitted to a parabola to interpolate the preferred size that minimises EC_50_ or optimises antigen sensitivity (See Fig. S6 for other receptors). **(I)** The fitted preferred CD2-CD58 size for each receptor. An F-test compared the preferred size of each receptor to the TCR. **(J)** The concentration of antigen required to induce 50% target cell killing (EC_50_) by primary human CD8+ T cells over the preferred CD2-CD58 size for the indicated antigen receptor (from panel I). The EC_50_ values are taken from Burton et al (4).

To test our hypothesis, we examined the effect of elongating CD2 by inserting increasing portions of the extracellular domain of the mucin CD43 (Fig. 3B). A key advantage of using a mucin is that it adopts an extended conformation and lacks tertiary structure (27), which enables precise nanometre control of extracellular size. As expected, wild-type CD2 increased TCR sensitivity while elongating it progressively decreased it (Fig. 3C). In contrast, elongating CD2 enhanced the sensitivity of all standard CARs (Fig. 3D-F). The sensitivity of the STAR format was optimised by wild-type CD2, consistent with its extracellular dimension being shared with the TCR (Fig. 3G).

Interestingly, the optimal CD2 spacer length varied from 8 to 50 amino acids of CD43 depending on the CAR hinge, suggesting that varying the hinge varies the size of the CAR/antigen complex. Since the dimensions of the relevant portions of CD43, CD2, and CD58 are known, we could estimate the CD2/CD58 complex size that optimised sensitivity for each antigen receptor (Fig. 3H, Fig. S6). This preferred CD2/CD58 size was similar for the TCR and STAR and progressively increased for CARs with the CD28, CD8a, and IgG1 hinges (Fig. 3I). Indeed, the ability of T cells expressing wild-type CD2 to kill target cells decreased for CARs that preferred larger CD2-CD58 sizes (Fig. 3J). Given that the preferred CD2-CD58 size reflects the size of the CAR/antigen complex, we used the known size of the pMHC antigen to estimate the size of each CAR finding that they all exceeded the extracellular size of the TCR (Table 1).

**Table 1:** Estimated Extracellular Sizes.

| Receptor | Preferred CD2/CD58 Size, nm (95% CI) | Receptor Size<sup>†</sup>, nm |
| --- | --- | --- |
| TCR | 14.8 | 7.8 |
| STAR | 15.0 (14.4 - 15.6) | 8.0 |
| CAR (CD28 H/TM-z) | 17.9 (17.3 - 18.2) | 10.9 |
| CAR (CD8a H/TM-z) | 20.2 (19.5 - 20.7) | 13.2 |
| CAR (IgG1 H/CD28-z) | 24.8 (24.2 - 25.6) | 17.8 |

<sup>†</sup>Determined by subtracting the 7 nm size of pMHC antigen from the preferred CD2/CD58 size

To assess whether engineering CD2 size can fully optimise CAR antigen sensitivity, we directly compared the antigen sensitivity of the TCR with the CD8a H/TM CAR, whose sensitivity was optimised by CD2-CD43(40). We found that CAR-T cells engineered with CD2-CD43(40) achieved the same optimal antigen sensitivity as TCR-T cells expressing wild-type CD2 when recognising antigen on Nalm6, U87, and T2 cell lines (Fig. 4A). Interestingly, this optimal sensitivity differed between the cell lines (optimal EC_50_ for Nalm6 *>* U87 *>* T2) and correlated with CD58 but not ICAM-1 surface expression on each cell line (Fig. 4B). We also confirmed that elongating CD2 increases the sensitivity and efficacy of primary CD4^+^ and CD8^+^ CAR-T cells (Fig. S7). Taken together, these results explain why different CARs exhibit differences in antigen sensitivity even when using the same signalling and antigen recognition domains, and demonstrates that engineering CAR-T cells with their preferred CD2/CD58 size optimises their antigen sensitivity to match that of the TCR.

**Figure 4:**
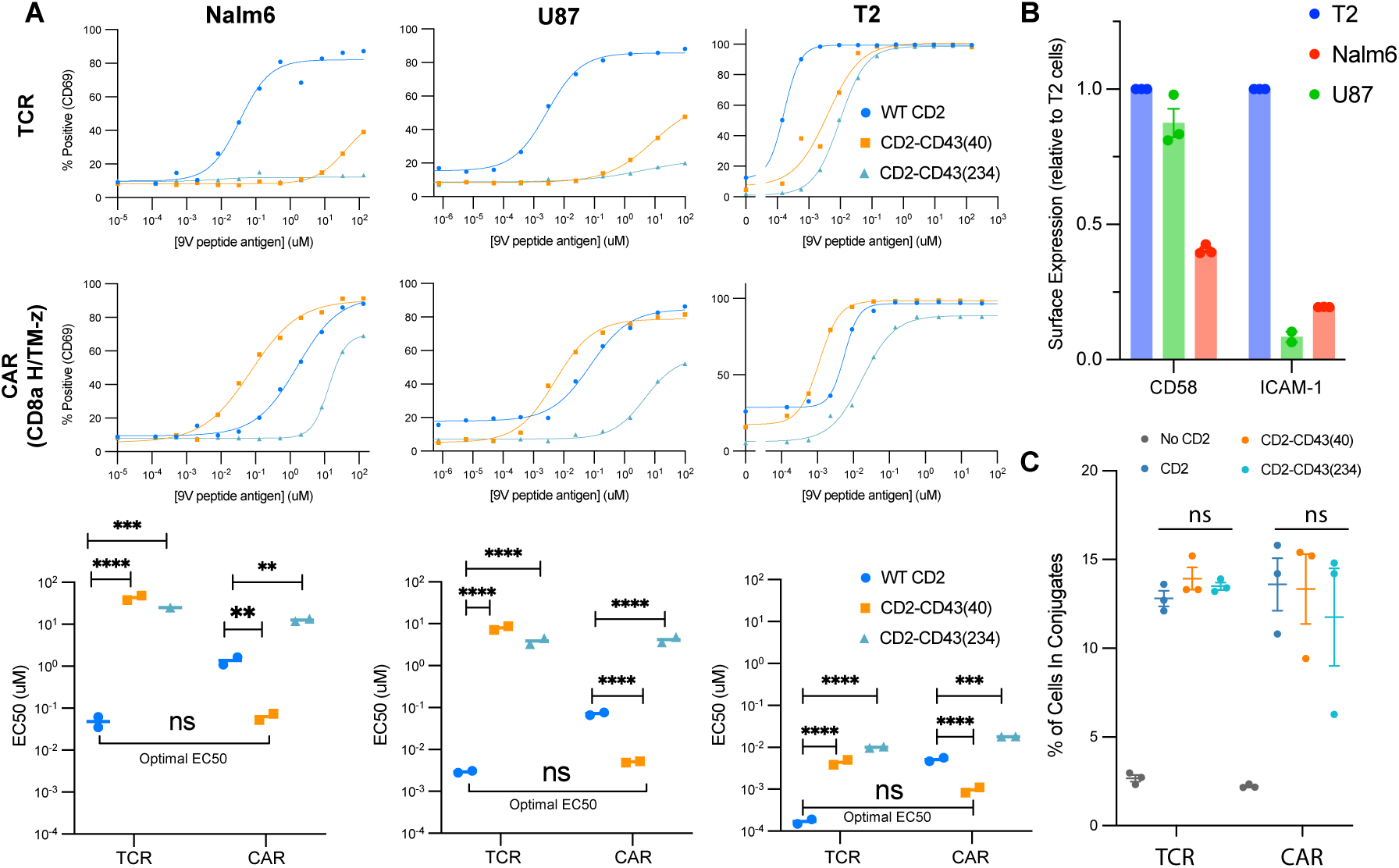
CAR-T cells engineered with optimally elongated CD2 achieve the same antigen sensitivity as TCR-T cells with wild-type CD2. **(A)** Jurkat T cells expressing the TCR or CAR engineered with the indicated CD2 recognising antigen on the B cell leukaemia Nalm6 cell line, glioblastoma U87 cell line, or the B cell hybridoma T2 cell line. Representative experiments (top) with summary EC_50_ across N=2 independent experiments (bottom). **(B)** Expression of the adhesion ligands CD58 and ICAM-1 on each cell line. **(C)** Percent of Jurkat T cells in conjugate with T2 cells. A one-way ANOVA with Holm-Sidak multiple comparison correction is used to determine p-values. Abbreviations: ** = p-value≤0.01, *** = p-value≤0.001, **** = p-value≤0.0001.

### The preferred CD2-CD58 size increases CAR/antigen binding and nanoscale co-localisation

To explore how size matching increases sensitivity, we first investigated how elongated CD2 impacts cell-cell conjugation. Unexpectedly, we found that all CD2 sizes tested mediated increased T cell/APC conjugation to a similar extent for both the TCR and the CAR (Fig. 4C). This indicates that changes in CD2/CD58 complex size do not affect overall cell/cell adhesion.

We next investigated whether CD2/CD58 size impacts the molecular organisation within the immune synapse using supported planar bilayers (SLBs) presenting antigen, adhesion ligands, and large glycocalyx components (28) (Fig. 5A). In the case of the TCR, CD58 and pMHC antigen co-localised within regions that excluded the glycocalyx components when T cells expressed wild-type CD2 but not elongated CD2 (Fig. 5B, top). In contrast, in the case of the CAR, the preferred elongated CD2-CD43(40) induced co-localisation of CD58 and pMHC antigen whereas segregation was observed with wild-type CD2 or excessively elongated CD2-CD43(234) (Fig. 5B, bottom). A mechanism by which colocalization within the synapse increases antigen sensitivity is through increased CAR/antigen engagement, since the preferred CD2/CD58 size would position the membranes at the optimal distance for the TCR or CAR to bind antigen. In support of this, elongating CD2 decreased antigen engagement by the TCR, while the preferred CD2-CD43(40) elongation increased antigen engagement by the CAR (Fig. 5C). For both the TCR and CAR, excessive elongation of CD2 using CD2-CD43(234) reduced antigen engagement relative to the preferred CD2 size.

**Figure 5:**
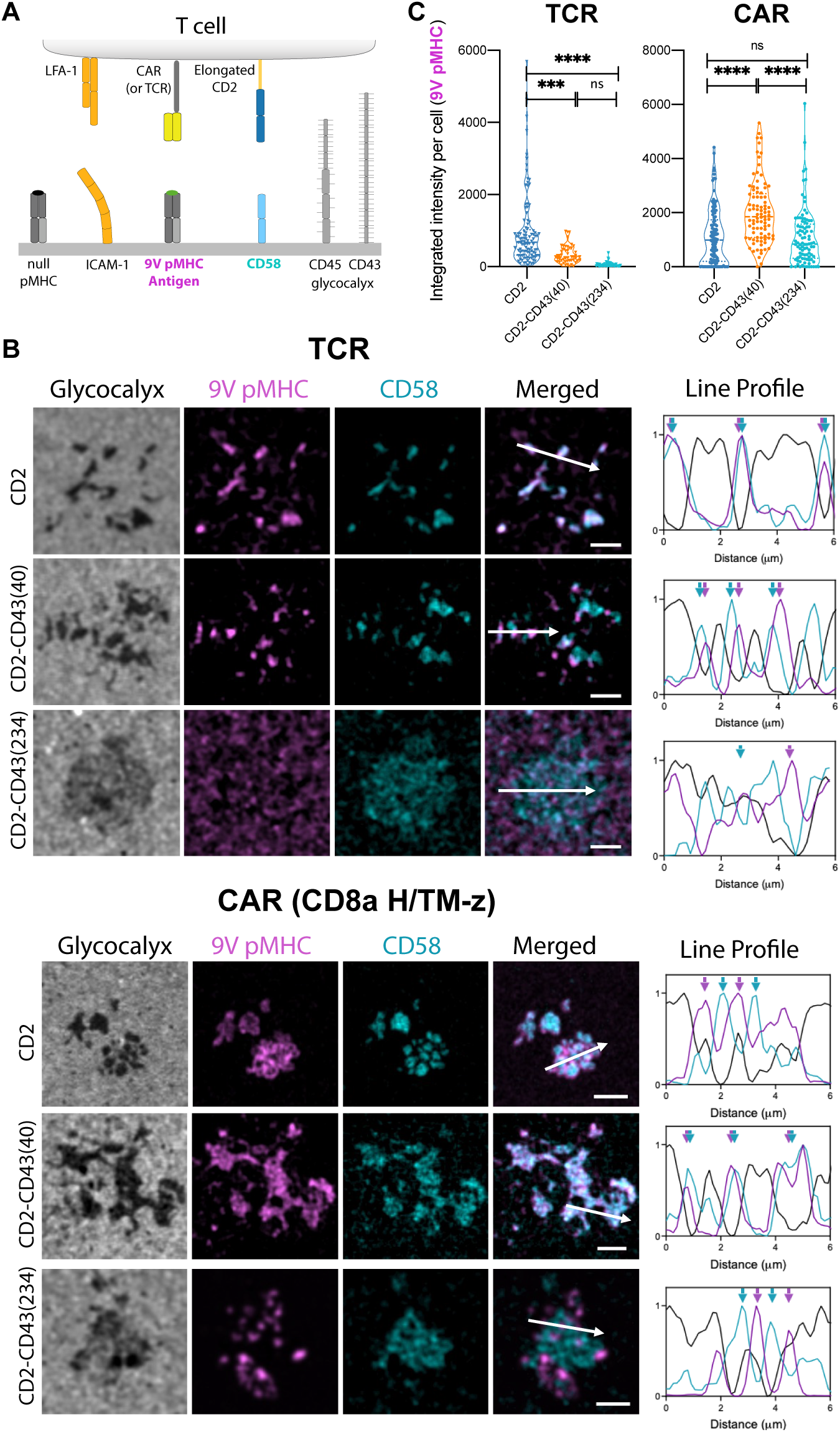
Matched CD2/CD58 and CAR/antigen size increase co-localisation and antigen engagement. **(A)** Supported planar bilayers 2.0 contain adhesion ligands (ICAM-1, CD58), antigen (9V pMHC), and large negatively charged glycocalyx components (CD45, CD43). **(B)** Jurkat T cells expressing a TCR (top) or a CAR (bottom) with the indicated CD2 molecule are imaged using TIRF microscopy (Scale bar is 2.5 *µ*m). Line profile intensities along the direction of the white arrow indicates CD58, pMHC antigen, and glycocalyx showing CD58/antigen co-localisation for the TCR only with wild-type CD2 and for the CAR only with optimally elongated CD2-CD43(40). **(C)** Integrated intensity of pMHC antigen under each cell for the indicated conditions. All p-values determined using a t-test with Holm-Sidak multiple comparison correction on log-transformed intensity values. Abbreviations: * = p-value≤0.05, ** = p-value≤0.01, *** = p-value≤0.001, **** = p-value≤0.0001.

Taken together with the functional data, these results are consistent with CD2/CD58 extracellular size determining the intermembrane distance (or contact height), which promotes antigen co-localisation and engagement only when the contact height matches the TCR or CAR/antigen size.

### Engineered contact heights for optimal CAR-T cell sensitivity reduce the potency of PD-1 co-inhibition

A potential consequence of optimising CAR sensitivity by artificially increasing the contact height through CD2 elongation is that it may reduce the activity of other co-signalling receptor/ligand interactions that rely on a physiological ∼14 nm height, such as the inhibitory PD-1/PD-L1 axis. Specifically, we reasoned that PD-1 would exhibit reduced inhibition of CARs within these elongated contacts unless PD-1 itself was artificially elongated to match the contact height (Fig. 6A). Strikingly, while elongating PD-1 diminished its capacity to inhibit the TCR, it enhanced its ability to inhibit the CAR (Fig. 6B). These results underscore the critical role of size matching for effective signal integration between surface receptors and reveal how a size mismatch can attenuate signal integration, which in the case of inhibitory receptors enables higher CAR-T cell sensitivity.

**Figure 6:**
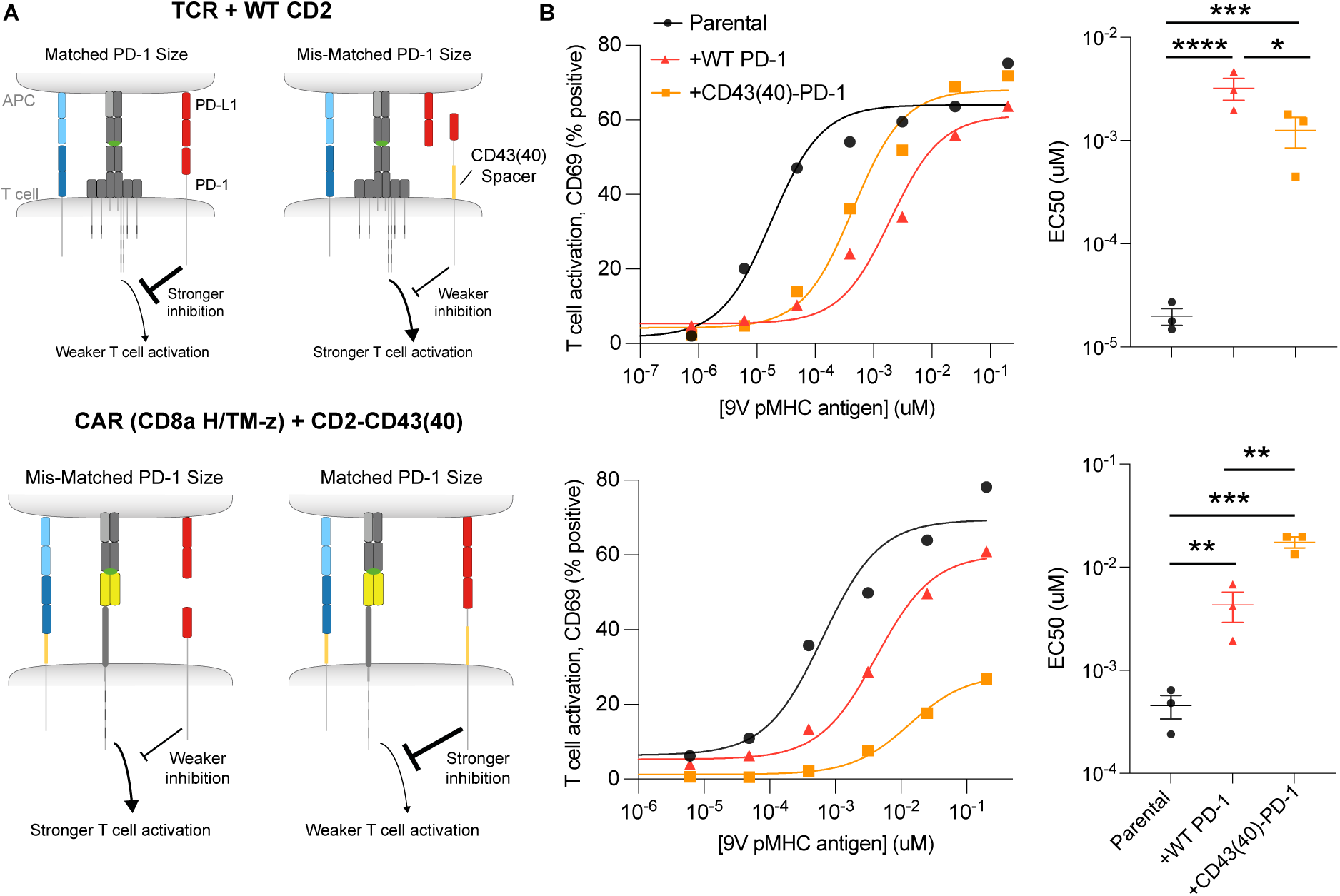
A mismatch in extracellular size diminishes the ability of PD-1 to inhibit CAR-T cell activation. **(A)** Cartoon to illustrate the hypothesis that elongating PD-1 will decrease its ability to inhibit the TCR (top) but can increase its ability to inhibit CARs (bottom). **(B)** Jurkat T cells expressing the TCR with wild-type CD2 (top) or the CAR with optimally elongated CD2-CD43(40) (bottom) transduced with PD-1 or elongated PD-1-CD43(40) are stimulated with CHO-K1 CombiCells presenting pMHC, CD58, and PD-L1. The CD43(40) spacer is selected to elongate PD-1 because elongating CD2 with this spacer optimised antigen sensitivity for the CAR. All p-values determined using a t-test with Holm-Sidak multiple comparison correction on log-transformed EC_50_ values. Abbreviations: * = p-value≤0.05, ** = p-value≤0.01, *** = p-value≤0.001, **** = p-value≤0.0001.

### The preferred CD2/CD58 size depends on the antigen epitope

The size of the CAR/antigen complex will depend not only on the CAR hinge but also on the antigen epitope distance from the target cell membrane. To explore this, we used scFvs known to target a membrane-proximal (m971) and membrane-distal (RFB4) epitopes of the large CD22 antigen (29) (Fig. 7A). As clinical trials targeting CD22 have used both the CD28 and CD8a hinges (30), we fused both scFvs to each hinge and expressed the resulting four CARs in T cells lacking CD2 or 7 different CD2 sizes resulting in 32 cell lines (Fig. S8A).

**Figure 7:**
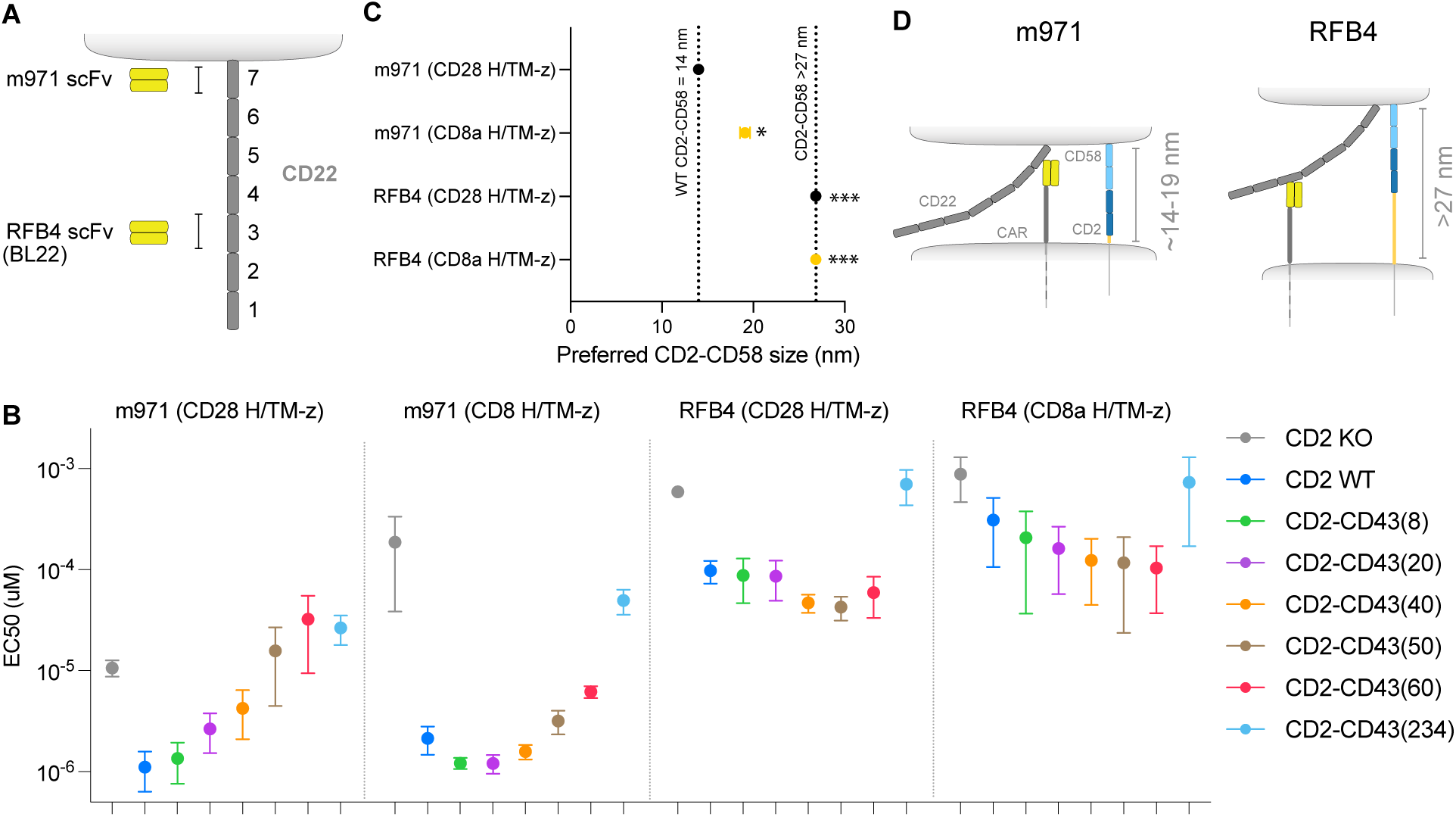
The CAR hinge and epitope impact the preferred size of CD2 required for optimising antigen sensitivity. **(A)** The m971 scFv binds a membrane-proximal and RFB4 binds a membrane-distal epitope of CD22. **(B)** T cell antigen sensitivity for the indicated CAR and CD2. Jurkat T cells with matched CAR and CD2 expression were co-cultured with CHO-K1 CombiCells presenting different concentrations of Spytag-CD22 with a fixed concentration of Spytag-CD58 (Fig. S8). The EC_50_ is the concentration of CD22 required for 50% of T cells to upregulate CD69 after 4 hours. **(C)** The preferred CD2-CD58 size for optimal antigen sensitivity. **(D)** Cartoon depicting the optimal intermembrane distance for the membrane-proximal (m971) and membrane-distal (RFB4) binders. An F-test compares each CAR to m971 (CD28 H/TM-z) in panel C. Abbreviations: * = p-value≤0.05, *** = p-value≤0.001.

We next co-cultured these cells with CHO-K1 CombiCells presenting CD58 and a range of CD22 surface densities (Fig. S8B), enabling accurate estimates of EC_50_ for CAR-T cell activation (Fig. 7B). The sensitivity of the membrane-proximal m971 CAR with the shorter CD28 hinge was most enhanced by wild-type CD2, while the sensitivity of the m971 CAR with the CD8a hinge was maximal with CD2 elongated by 8-20 amino acids of CD43. In contrast, the sensitivity of the membrane-distal RFB4 CAR fused to either hinge increased with elongation of CD2 by up to 50-60 amino acids, but these increases were modest and appeared to saturate. The preferred CD2/CD58 complex size for each CAR ranged from 14 to ≳27 nm (Fig. S9, Fig. 7C). Since CD22 forms a rod-like conformation that extends ∼30 nm (31), the extracellular region of CD22 likely adopts a tilted orientation when engaged by m971 CARs to enable the membranes to form a sufficiently close contact (14-16 nm apart, Fig. 7D).

## Discussion

Our results show that a mismatch in size between CAR/antigen and CD2/CD58 complexes contributes to the low sensitivity of CAR-T cells and that engineering more compact CARs or engineering elongations of CD2 using portions of the CD43 mucin can fully restore CAR-T cell sensitivity. These results have important implications for tuning CAR-T cell sensitivity and for our basic understanding of the TCR and CD2.

There is increasing evidence that CD2 is crucial for enabling T cells to recognise antigen on cancerous and infected cells (21–25). A current model for how CD2 increases T cell antigen recognition is that it facilitates the formation of micovillar contacts that enable the TCR to bind pMHC on target cells (28). Consistent with this hypothesis, we have found that wild-type and elongated CD2 facilitated microvillar contacts that excluded glycocalyx components for both the TCR and CAR. Additionally, we now report that efficient antigen engagement within these contacts requires matched sizes: while elongated CD2-CD43(40) continued to form microvillar contacts it reduced antigen engagement for the TCR yet increased it for the CAR relative to wild-type CD2. This suggests that nanoscale elongations of CD2 disrupt its ability to generate a contact height of ∼14 nm required for TCR to bind pMHC (32–35). Thus, the CD2/CD58 adhesion interaction facilitates microvillar contacts at precise contact heights that can be manipulated through CD2 elongations to potentially match a wide array of antigen-receptor/antigen sizes.

While STARs/HITs can display higher antigen sensitivity compared to standard CARs, it has been unclear whether this is a result of changes to intracellular signalling or extracellular size (4, 10–13). To understand their relative contributions, we generated a Compact CAR that uses standard CAR signalling (28z) but matches the extracellular size of the TCR (2 Ig domains). We note that standard CARs form homodimers because they use hinges (CD28, CD8a, IgG) that mediate homodimersation in their native proteins (36–38). Therefore, standard CARs likely contain two copies of the 28z signalling chains like Compact CARs. As a result, the much higher sensitivity of the Compact CAR and its ability to exploit wild-type CD2/CD58 adhesion is likely a result of its matched extracellular dimensions rather than differences in signalling with standard CARs. Indeed, we found that we can increase the sensitivity of standard CARs by elongating CD2 or reducing the size of their hinge. For example, we found that pMHC and CD22-targeting CARs with a CD28 hinge preferred a smaller CD2-CD58 size compared to CARs relying on a CD8a hinge. This implies that the CD28 hinge is shorter than the CD8a hinge, and this may explain why Yescarta (with a CD28 hinge) achieves higher sensitivity than Kymriah (with a CD8a hinge) when recognising the same CD19 antigen (5, 15).

The hypothesis that extracellular size is crucial for receptor/ligand interactions at cellular interfaces was first proposed in the early 1990s (39–41). The experimental evidence supporting this is based on the demonstration that elongation of surface molecules impairs their function. For instance, previous work has shown that using large globular domains to elongate the TCR or CAR interaction with their respective antigens or the CD2 and PD-1 receptors with their ligands results in diminished function (17, 42–45). However, these findings could be attributed to size-independent mechanisms, whereby elongation of surface receptors changes their structure or interactions with other molecules for example. In the present study, we have introduced nanoscale elongations of CD2 and PD-1 using portions of the CD43 mucin confirming reduced activity when acting on the TCR but, importantly, we show that the same molecules exhibit increased activity when acting on CARs. Collectively, these results now demonstrate that nanoscale changes in receptor/ligand sizes can profoundly impact their function and offers a new way to precisely control extracellular sizes.

Taken together, our work demonstrates that diverse CAR/antigen sizes can be generated by chimeric receptors with different extracellular architectures targeting different antigens, and different epitopes on the same antigen. As a result, each CAR/antigen pair prefers a different CD2/CD58 size for optimising their antigen sensitivity. We hypothesised that elongating CD2 to form artificial contact heights that exceed 14 nm would disrupt other co-signalling receptor/ligand interactions. Consistent with this, we found that the potency of wild-type PD-1/PD-L1 co-inhibition is reduced unless it is elongated to match the CAR/antigen size. Given that many co-stimulation and co-inhibition receptors are proposed to span the same 14 nm intermembrane distance when interacting with their ligands (46), our work suggests a general paradigm where forming artificially spaced contacts can be used to disrupt native receptor/ligand interactions and selectively introduce interactions by size-based re-engineering. These artificially large contacts may also reduce segregation from the phosphatase CD45, which is critical for antigen receptor triggering by the kinetic-segregation mechanism (17, 46–48). This may explain why elongating CD2 could not fully rescue CD22-targeting CARs that rely on the membrane-distal RFB4 scFv where the CAR/antigen size may be too large to mediate effective CD45 segregation. Overall, our results demonstrate that antigen sensitivity and signal integration in CAR-T cells can be optimised by engineering the extracellular sizes of CAR/antigen and/or co-signalling receptor/ligand complexes.

## Open access

This research was funded in whole, or in part, by the Wellcome Trust [207537/Z/17/Z]. For the purpose of Open Access, the author has applied a CC BY public copyright licence to any Author Accepted Manuscript version arising from this submission.

## Author contributions

Conceptualization (OD), Data Curation (JB, JAS, VA, EJ, MIB), Formal Analysis (JB, JAS, VA, EJ, MIB), Funding Acquisition (OD), Investigation (JB, JAS, VA, EJ, MIB, JCC, SJD, TRW, PAvdM, OD), Methodology (JB, JAS, VA, EJ, MIB, SBE), Project Administration (OD), Supervision (SJD, PAvdM, OD), Visualization (JB, JAS, VA, EJ, OD), Writing – Original Draft (OD, PAvdM), Writing – Review & Editing (JB, JAS, VA, EJ, MIB, SBE, JCC, SJD, TRW, PAvdM, OD)

## Acknowledgements

We thank Marion H. Brown and Sophie Heese for helpful discussion. We thank the Don Mason Flow Cytometry Facility.

## Materials & Methods

### Peptides

The NY-ESO-1_157-165_ derived 9V peptide (SLLMWITQV) was synthesised to *>* 95% purity (Peptide Protein Research, UK or GenScript, UK).

### Proteins

Production of Spytag-pMHC: HLA-A*02:01 heavy chain (UniProt residues 25–298) with a C-terminal Spytag003 and β_2_-microglobulin were expressed as inclusion bodies in *E.coli*, refolded *in vitro* as described in (49) together with the 9V NY-ESO-1 peptide, and purified using size-exclusion chromatography on a Superdex S75 column (GE Healthcare, USA) in HBS-EP buffer (10 mM M HEPES pH 7.4, 150 mM NaCl, 3 mM EDTA, 0.005% v/v Tween-20).

Production of Spytag-CD58/PD-L1/CD22: Expi293™ cells (ThermoFisher Scientific, A14527) were grown in Expi293™ Expression Medium (ThermoFisher Scientific, A1435101) in a 37°C incubator with 8% CO2 on a shaking platform at 130 rpm. Cells were passaged every 2–3 days with the suspension volume always kept below 33.3% of the total flask capacity. The cell density was kept between 0.5 and 3 million per ml. Before transfection cells were counted to check that cell viability was above 95%, and the density was adjusted to 3.0 million per ml. For 100 ml transfection, 320 µl ExpiFectamine™ 293 Transfection reagent (ThermoFisher Scientific, A14524) was mixed with 6 ml Opti-MEM (ThermoFisher Scientific, 31985062) for 5 min. During this incubation, 100 µg of expression plasmid was mixed with 6 ml Opti-MEM. The DNA was then mixed with the ExpiFectamine™ and incubated for 15 min before being added to the cell culture. One day after transfection 600 µl of enhancer 1 and 6 ml of enhancer 2 was added to the culture flask. The culture was returned to the shaking incubator for 4-5 days for protein expression to take place.

Cells were harvested by centrifugation and the supernatant collected and filtered through a 0.22 µm filter. Imidazole was added to a final concentration of 1 mM and PMSF added to a final concentration of 1 mM; 2 ml of Ni-NTA Agarose (Qiagen, 30310) was added per 50 ml of supernatant and the mix was left on a rolling platform at 4°C overnight. The mix was poured through a gravity flow column to collect the Ni-NTA Agarose. The Ni-NTA Agarose was washed three times with 10 ml of wash buffer (50 mM NaH2PO4, 300 mM NaCl, and 5 mM imidazole at pH 8). The protein was eluted with 15 ml of elution buffer (50 mM NaH2PO4, 300 mM NaCl, and 250 mM imidazole at pH 8). The protein was concentrated, and buffer exchanged into size exclusion buffer (25 mM NaH2PO4 and 150 mM NaCl at pH 7.5) using a protein concentrator with a 10,000 molecular weight cut-off. The protein was concentrated down to 500 µl and loaded onto a Superdex 200 10/300 GL (Cytiva, 17-5175-01) size exclusion column. Fractions corresponding to the desired peak were pooled and frozen at –80°C. Samples from all observed peaks were analysed on a reducing SDS–PAGE gel.

For purified Spytag-CD22, SUMO was used to stabilise the protein during production and therefore the HRV 3C Protease Solution Kit was used for SUMO removal (Pierce™, 88946). HRV protease was added to the purified protein at a pre-determined optimum ratio for full cleavage of the HRV site. The mixture was left overnight for full cleave to occur and then 1 ml of Glutathione Agarose (Pierce™, 16100) added for 4 hours to remove the protease. The solution was run through a gravity flow column to collect to SUMO plus protein of interest mixture. This was then added to 1 ml of Ni-NTA Agarose (Qiagen, 30310) and left on a rolling platform at 4°C overnight. The mix was poured through a gravity flow column to collect the Ni-NTA Agarose. The Ni-NTA Agarose was washed once with 10 ml of wash buffer (50 mM NaH2PO4, 300 mM NaCl, and 5 mM imidazole at pH 8). The protein was eluted with 15 ml of elution buffer (50 mM NaH2PO4, 300 mM NaCl, and 250 mM imidazole at pH 8). The protein was concentrated, and buffer exchanged into size exclusion buffer (25 mM NaH2PO4 and 150 mM NaCl at pH 7.5) using a protein concentrator with a 10,000 molecular weight cut-off and frozen in suitable aliquots at –70°C.

Spytag CD58, PD-L1, and CD22 contained the full extracellular domain fused to a c-terminal Spytag003 (RGVPHIVMVDAYKRYK) followed by a Histag for purification (HHHHHH).

All purified proteins were stored at -70°C until use.

### Production of primary human CD8+ TCR/CAR-T cells

HEK 293T cells were seeded in DMEM supplemented with 10% FBS and 1% penicilin/streptomycin in 6-well plates to reach 60–80% confluency on the following day. Cells were transfected with 0.25 pRSV-Rev (Addgene, 12253), 0.53 *µ*g pMDLg/pRRE (Addgene, 12251), 0.35 *µ*g pMD2.G (Addgene, 12259), and 0.8 µg of transfer plasmid using 5.8 X-tremeGENE HP (Roche). Media was replaced after 16 hours and supernatant harvested after a further 24 hours by filtering through a 0.45 *µ*m cellulose acetate filter. Supernatant from one well of a 6-well plate was used to transduce 1 million T cells.

Human CD8+ T cells were isolated from leukocyte cones purchased from the National Health Service’s (UK) Blood and Transplantation service. This project has been approved by the Medical Sciences Inter-Divisional Research Ethics Committee of the University of Oxford (R51997/RE001), and all leukocyte cones were anonymised by the NHS before purchase. As a result of using anonymised leukocyte cones, we did not apply any exclusion or inclusion criteria, the attrition rate was nill, the sex, gender, age, and weight of donors were unknown and hence were not variables in the present study. As donors were simply used as a source of T cells, we did not have subject groups and therefore, did not require randomisation of subject groups and hence blinding was not required. The smallest effective size that we aimed to resolve was a 3-fold change in sensitivity (a difference of 0.47 on log-transformed EC50 values) and a power calculation shows that this can be be resolved with a power of 80% (alpha at 0.05) using three samples in each group. Therefore, all experiments will be repeated with 3 independent donors.

Isolation was performed using negative selection. Briefly, blood samples were incubated with Rosette-Sep Human CD8+ enrichment cocktail (Stemcell) at 150*µ*l/ml for 20 minutes. This was followed by a 3.1 fold dilution with PBS before layering on Ficoll Paque Plus (GE) at a 0.8:1.0 ficoll to sample ratio. Ficoll-Sample preparation was spun at 1200g for 20 minutes at room temperature. Buffy coats were collected, washed and isolated cells counted. Cells were resuspended in complete RMPI (RPMI supplemented with 10% v/v FBS, 100 penicillin, 100 streptomycin) with 50U of IL-2 (PeproTech) and CD3/CD28 Human T-activator Dynabeads (Thermo Fisher) at a 1:1 bead to cell ratio. At all times isolated human CD8+ T cells were cultured at 37 and 5% CO2. T cells at 1 million cells in 1 ml of media were subsequently transduced on the following day using lentivirus encoding for the desired antigen receptor. On days 2 and 4 post-transduction, 1ml of media was exchanged and IL-2 was added to a final concentration of 50U. Dynabeads were magnetically removed on day 5 post-transduction. When using the TCR, T cells were further cultured at a density of 1 million/ml and supplemented with 50U IL-2 every other day. When using CARs, T cells were further cultured at a density of 0.5 million/ml and supplemented with 100U IL-2 every other day. T cells were used between 10 and 16 days after transduction.

### Production of Jurkat TCR/CAR-T cells

The previously described TRAC^(−*/*−)^TRBC^(−*/*−)^ E6.1 Jurkat T cells (50) were used as the parental Jurkat line to knockout CD2.

To produce 1 million T cells, 2 million cells are electroporated with guides and Cas9 in the following protocol. The Cas9 and guide RNAs, which are TrueGuide (ThermoFisher) synthetic modified guide RNAs with the following sequences: CAAGGCACCCCAGGTTTCCA, CAAAGAGATTACGAATGCCT, CTTG-TAGATATCCTGATCAT and, GCATCTGAAGACCGATGATC, are mixed: 3.6 µL of OptiMEM (Gibco),

3.4 µL of TrueCut Cas9 V2 (ThermoFisher), 0.75 uL of gRNA from 100 µM stocks and incubated at room temperature for at least 10 minutes. The cells are washed 3x in OptiMEM and resuspended in 50 µL of OptiMEM per million cells. 2 million cells (100 µL) are mixed with the Cas9/gRNA preperation and transferred to a electroporation cuvette (Cuvette Plus, 2 mm gap, BTX) and electroporated (300V, 2 mesc, ECM 830 Square Wave Electroporation System), then immediately transferred to 1 mL of prewarmed cell culture media. Following cell sorting, we termed these CD2 KO Jurkat T cells.

CD2 KO Jurkat T cells were transduced with the desired antigen receptor followed by CD2 WT or the desired elongated CD2 molecules. In the case of the TCR and the CD8a hinge CAR we tested a panel of smaller sizes first and then repeated the experiments with a second panel that included larger CD2 sizes (Fig. 3C,D). To investigate PD-1, these cells were further transduced with PD-1 wild-type or elongated PD-1. Jurkat T cells were sorted for matched expression of introduced surface proteins.

### Production of primary human CD8+ CAR-T cells expressing elongated CD2

To produce primary human CD8+ CAR-T cells expressing elongated CD2, the CRISPR protocol above for CD2 KO was performed following negative selection of primary human CD8+ or CD4+ T cells and before addition of IL-2 and CD3/CD28 Dynabeads (see protocol above for ‘Production of primary human CD8+ TCR/CAR-T cells’). The lentivirus used encodes the CAR (CD8a H/TM-z) and either CD2 WT or CD2-CD43(40).

### Co-culture of U87s with Jurkat or primary human T cells

On day 0 25,000 U87 cells were seeded in a flat-bottom TC-treated 96 well plate. On day 1 the media was carefully removed from these cells and they were incubated with peptide at the indicated concentrations diluted in DMEM supplemented with 10% FBS for 1 hour at 37°C. The peptide containing media was then removed and 50,000 T cells in 100 uL of RPMI (supplemented with 10% FBS) and the co-culture spun for 2 minutes at 50 xg. For Jurkat T cells, cells were then incubated for 4 hours at 37°C. For primary human T cells, cells were incubated for 20 hours at 37°C. After this time a fraction of the supernatant was removed and used as described in the section on ELISAs. The remaining cells were detached by pipetting and subsequently stained in PBS 1% BSA for CD45 (HI30, 1:200), CD69 (FN50, 1:200) and, in the case of primary T cells, 41BB (4B4-1, 1:200). Stained cells were analysed immediately or fixed with PBS 1% formaldehyde and analysed on the following day. T cells were discriminated from U87 cells by CD45 expression and singlets identified on the basis of size; subsequent analysis was performed on this population.

### Co-culture assay with T2 cells and Jurkat T cells

T2 cells were stained with 5 µM Tag-It Violet (BioLegend) following the manufacturer’s protocol and then 60000 cells were seeded in a volume of 100 µL per well in a V-bottom 96 well tissue culture plate. T2 cells were then incubated with 100 µL of peptide dilution prepared to the desired concentration in complete RPMI for 1 hour at 37 °C. T2 cells were then washed, resuspended in 100 µl of complete RPMI and transferred to a flat-bottom 96 well tissue culture plate.

T cells were counted and re-suspended in fresh media such that there were 30000 cells per 100 µl. This volume was then added to the T2 cells transferred previously. Plates were then spun at 50 xg for 2 minutes and incubated for 8 hours at 37 °C. Cells were subsequently harvested as above. T cells were discriminated from T2 cells by the absence of Tag-It Violet stain. Single T cells were identified on the basis of size and subsequent analysis performed on this population.

### Co-culture assay with Nalm6 cells and Jurkat T cells

On the day of the experiment 50000 NALM6 cells were seeded in 100 µL per well in a V-bottom 96 well tissue culture plate. T2 cells were then incubated with 100 µL of peptide dilution prepared to the desired concentration in complete RPMI for 1 hour at 37 °C. T2 cells were then washed, resuspended in 100 µl of complete RPMI and transferred to a flat-bottom 96 well tissue culture plate. T cells were counted, resuspended and added to the NALM6 cells such that there were 50,000 T cells per well. Plates were then spun at 50 xg for 2 minutes and incubated for 6 hours at 37 °C. Cells were subsequently harvested as above.

### Co-culture assays with CombiCells

50,000 (Fig. 1E, Fig. S3, Fig. 6) or 25,000 (Fig. 7, S8) CHO-K1 CombiCells cells (5) were seeded in flat-bottom 96-well plates and incubated overnight at 37 degrees, 10% CO2. The antigen (9V pMHC or CD22) and relevant ligands (CD58, PD-L1) fused to Spytag003 were diluted to the required concentration in DMEM 10% FBS 1% Penicillin-Streptomycin. The media was removed and the prepared ligands added to the cells for an incubation of 40 min at 37 degrees, 10% CO2. The media (with any remaining ligands) was removed and the cells washed in 100 ul of DMEM media after which the media was removed again. The indicated T cells were added to the CHO-K1 CombiCells coupled to the indicated ligands at an E:T ratio of 1:2 for a 6 hr (50,000 T cells/well, Fig. 1E, Fig. S3, Fig. 6) or 1:1 for a 4 hr (50,000 T cells/well, Fig. 7, S8) co-culture assay at 37 degrees, 5%CO2. The cells were spun down for 1 min at 300 rpm at the start of the co-culture. All CombiCells relied on a titration of antigen with a fixed concentration of CD58 and/or PD-L1 (0.1 *µ*M).

### Cytotoxicity assay

Cytotoxicity was measured as LDH release using CyQUANT™ LDH Cytotoxicity Assay kits (ThermoFisher, C20300) following the manufacturers protocol. 45 minutes prior to the end of the assay the appropriate volume of 10X lysis buffer was added to control wells containing only effector or target cells and the same volume of sterile water was added to volume correction wells. Prior to harvesting plates were spun briefly at 5 xg for 3 minutes and then 50 µL of supernatant was carefully removed for the assay.

### Flow cytometry assay

At the end of the stimulation assay, the supernatant was carefully removed and saved for ELISA analysis. The cells were then aspirated and transferred to a v-bottom plate and washed once in 200*µ*l PBS 1% BSA (500g, 4C, 5 minutes). Antibodies against T cell activation markers were diluted in PBS 1% BSA at a 1:200 dilution. An anti-CD45 antibody was used to selectively stain T cells and distinguish them from CHO cells during flow cytometry analysis. To detect TCR/CAR expression fluorescently-conjugated peptide-MHC tetramers were added to the staining antibodies at a 1:1000 dilution. A viability dye was also added at a dilution if 1:2500 to distinguish live cells from dead cells. 50*µ*l of this staining solution was to the cells, before incubating them for 20 minutes at 4C in the dark. The cells were washed twice in PBS, and resuspended in 75*µ*l PBS, before running on a flow cytometer. Flow cytometry data was analysed using FlowJo (BD Biosciences, RRID:SCR-008520).

### Cytokine assay

IL-2 Human uncoated ELISA kit, TNF-*α* Human uncoated ELISA kit, or IFN-*γ* Human uncoated ELISA kit and Nunc MaxiSorp 96-well plates were used according to the manufacturer’s instructions. The supernatant from stimulation assays were diluted using an empirically determined factor prior to ELISAs to ensure that measurements were within the linear range of the standard curve. The absorbance at 450 nm and 570nm were measured using a SpectraMax M5 plate reader (Molecular Devices). Supernatants were either used immediately for ELISAs post-harvesting or stored at -20 °C for up-to 2 weeks.

### Conjugation assay

Jurkat T cells were tagged with CellTrace violet (ThermoFisher) and target cells were tagged with CellTrace far red (ThermoFisher) following the manufacturer’s instructions. The labelled cells were then mixed in equal quantities, centrifuged at 50 xg for 5 minutes and then incubated for 1 hour at 37°C. Cells were then gently resuspended and analysed by flow cytometry. Gates were set using single stained cells and the % of double positive cells is reported as the % conjugate formation.

### Airyscan confocal imaging of T cells on supported lipid bilayers

‘Second-generation’ glass-supported lipid bilayers (SLBs) were prepared as described previously (28). This SLB system contains physiological levels of pMHC^null^, pMHC^9V^, CD58, ICAM-1, and a glycocalyx mimetic (CD43 and CD45RABC) found on various antigen presenting cells. Close contacts formed by T cells (i.e. where TCR and CD2 are engaged), are observed as areas of exclusion of the glycocalyx mimetic (seen as a drop in glycocalyx fluorescence intensity). pMHC^9V^ was labelled with Alexa Fluor-488 (pMHC9V-AF488), CD58 with Alexa Flour-647 (CD58-AF647) and the glycocalyx (i.e., CD43 and CD45RABC) with Alexa Fluor-555. All other proteins were unlabelled. SLBs were produced using vesicle fusion. Lipid mixture consisting of 98% (mol%) POPC (Avanti Polar Lipids) and 2% (mol%) DGS-NTA-Ni2+ (NiNTA; Avanti Polar Lipids) in chloroform were mixed in a cleaned glass vial and dried under a stream of nitrogen. 2% NiNTA was chosen to minimise unwanted interactions of the lipid with biological material (e.g., non-tag histidine within proteins and negatively charged sugars on cells) whilst allowing physiological densities of multiple his-tagged proteins to be attached to the surface. The dried lipid mix was resuspended in 0.22 µm filtered PBS at 1 mg/ml, vortexed, and sonicated (30 min on ice with a tip sonicator, or until transparent in an ultrasonicator bath) to produce small unilamellar vesicles (SUVs). Glass coverslips (25 mm, thickness no. 1.5; VWR) were cleaned at least 2 h in 3:1 sulfuric acid/hydrogen peroxide at room temperature, rinsed in MQ water, and plasma cleaned for 1 min (oxygen plasma) or 20 min (argon plasma). CultureWell 50-well silicon covers (Grace Bio-Labs) were cut and placed on the washed coverslips (max 4/coverslip). SUVs were added to each well at a final concentration of 0.5 mg/ml (10 µl total volume) and left for 1 hour at room temperature. Wells were washed at least five times by removing and adding PBS to each well. The amount of liquid in each well was adjusted to be level with the well edge before adding 5 µl of proteins mixed at the desired concentrations (see (28) for concentrations). This was done to ensure reproducible incubation concentrations. Protein mixes were incubated with the bilayer for an hour at room temperature and washed ten times in pre-warmed (37°C) PBS + 2mM MgS0_4_ immediately before use.

5x10^4^ of the indicated cell line were resuspended in pre-warmed (37°C) PBS + 2mM MgS0_4_ and added to a washed SLB for 10-minutes at 37°C. 10 µl of the imaging buffer from the SLBs was then removed and replaced with 4% PFA + 0.25% glutaraldehyde for 15 minutes at room temperature prior to imaging. Imaging was performed on an LSM 880 scanning confocal with AiryScan module. The proteins on the SLB, i.e., pMHC9V-AF488, Glycocalyx-AF555, and CD58-AF647, were imaged using a NA1.4 63x oil immersion objective. Line scan averaging was set to a maximum of 16. Argon 488 nm, DPSS 561 nm and He-Ne 633 nm laser power set at appropriate levels to avoid bleaching and oversaturation. For measurements, the pinhole was set to 1 AU and a 488/594/633 MBS was used.

Cells were manually segmented and IRM contact area was quantified using Yen thresholding and the Analyze particle feature in FIJI.

### Data analysis

The dose-response curves were fitted to a Hill function in Prism (GraphPad Software, RRID:SCR-002798) to determine the EC_50_ (concentration of ligand producing 50% of the maximum response) and the E_max_ (maximum response at high ligand concentrations). When analysing cytokines where large differences in E_max_ were often observed, we calculated EC_15_ (instead of EC_50_) as the concentration required to elicit 15% of the largest E_max_ in the dataset. As a result, EC_15_ measures the concentration of antigen require to elicit the same concentration of cytokine across different antigen receptors/conditions, whereas the EC_50_ for each antigen receptor corresponds to the concentration of cytokine required to elicit 50% of that antigen receptors maximum, and hence EC_50_ can correspond to different levels of cytokine. All statistical analyses were carried out using Prism as described in figure captions.

### Estimating the preferred CD2-CD58 size

The size of the CD2/CD58 complex was estimated to be 14 nm based on X-ray crystallography and modelling (33). Available electron microscopy of the mucin CD43 reveals an extended conformation whereby each amino acid contributes on average 0.2 nm (27). Lastly, the short flexible linker between CD2 and CD43 (GSSSS) is estimated to contribute 0.85 nm, based on assuming a persistence length of 0.38 nm using the worm-like-chain model (i.e. 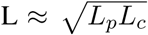 where *L_p_* = 0.38 nm for a flexible polypeptide and *L_c_* is the contour length of 5 aa x 0.38 nm). Therefore, CD2-CD43(40)-CD58 is estimated to be 22.8 nm.

We next fit plots of log-transformed EC_50_ over the estimated CD2-CD58 size using a quadratic equation

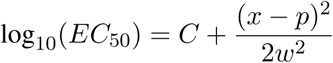

where *p* is the preferred CD2-CD58 size and w is the width. In the case of the CD28, CD8a, and IgG1 hinge CARs, we fitted both *p* and *w* because a complete parabola could be observed. However, this was not possible for the TCR and STAR because wild-type CD2 appeared to optimise sensitivity. Therefore, in the case of the TCR, we fixed p to the wild-type CD2-CD58 size and fit only for *w* because structural studies have shown matched dimensions (35). In the case of the STAR, we fixed *w* to the value obtained for the TCR and fitted *p*.

### Sequences

The sequences of surface molecules used in the present study are summarised below and in Fig. S1.

D52N scFv (VH-Linker-VL):

MAEVQLLESGGGLVQPGGSLRLSCAASGFTFSTYQMSWVRQAPGKGLEWVSGIVSSGGSTAYADS VKGRFTISRDNSKNTLYLQMNSLRAEDTAVYYCAGELLPYYGMDVWGQGTTVTVSS AKTTPKLEEGEFSEARVQSELTQPRSVSGSPGQSVTISCTGTERDVGGYNYVSWYQQHPGKAPKLII HNVIERSSGVPDRFSGSKSGNTASLTISGLQAEDEADYYCWSFAGGYYVFGTGTDVTVLGQPKANPT VDPDYKDDDDK

IgG1-CH1:

ASTKGPSVFPLAPSSKSTSGGTAALGCLVKDYFPEPVTVSWNSGALTSGVHTFPAVLQSSGLYSLSSVVTVPSSSLGTQTYICNVNHKPSNTKVDKRVEPKSC

IgG1-CL:

GQPKANPTVTLFPPSSEELQANKATLVCLISDFYPGAVTVAWKADGSPVKAGVETTKPSKQSNNKYAAS SYLSLTPEQWKSHRSYSCQVTHEGSTVEKTVAPTECS

HNG Spacer:

DPAEPKSPDKTHTCPPCP

Furin-P2A:

GSRAKRSGSGATNFSLLKQAGDVEENPGP

### Antibodies

CD45: Clone HI30 BV421 Biolegend RRID:AB2561357

CD3: Clone OKT3 488 Biolegend RRID:AB571877; Clone UCHT1 BV421 Biolegend RRID: AB10962690;

4-1BB: Clone 4B4-1 AF647 Biolegend RRID:AB2566258; Clone 4B4-1 BV421 Biolegend RRID: AB2563830;

CD69: FN50 AF647 Biolegend RRID:AB528871; Clone FN50 A488 Biolegend RRID: AB528869; Clone FN50 BUV395 BD Biosciences RRID: AB2738770;

CD2: Clone TS1/8 PE Biolegend RRID:AB314758; Clone TS1/8 BV421 Biolegend RRID: AB2561832;

TCR V13.1: Clone H131 APC Biolegend RRID: AB2728348;

CD19: Clone H1B19 BV421 Biolegend RRID: AB11142678;

PD-1: Clone EH12.2H7 FITC Biolegend RRID: AB940479;

PDL1: Clone MIH3 PE Biolegend RRID: AB2734439;

HLA A,B,C: Clone W6/32 A488 Biolegend RRID: AB493134;

HA Epitope: Clone 16B12 PE Biolegend RRID: AB2629623;

CD2: Clone TS1/8 FITC Biolegend RRID:AB2876780;

CD22: Clone HIB22 BV421 Biolegend RRID:AB2721514

CD58: Clone TS2/9 PE Biolegend RRID:AB1186063

CD58: Clone TS2/9 APC Biolegend RRID:AB2650886

PD-1: Clone EH12.2H7 PE Biolegend RRID:AB940481

## Data Availability

This study includes no data deposited in external repositories.

## Supplementary Information

**Figure S1:**
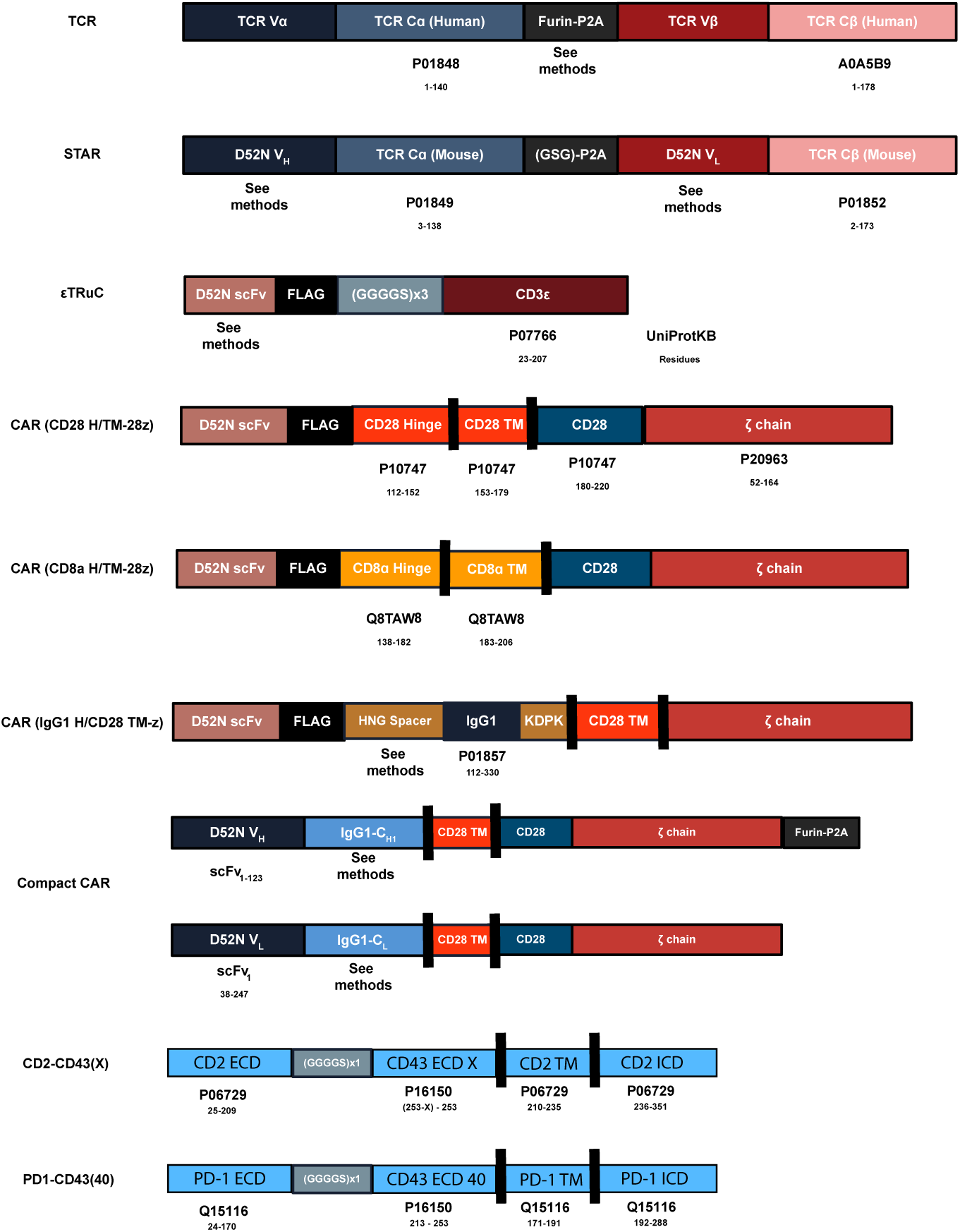
Architecture and sequences of surface molecules used in the present study (also see Methods).

**Figure S2:**
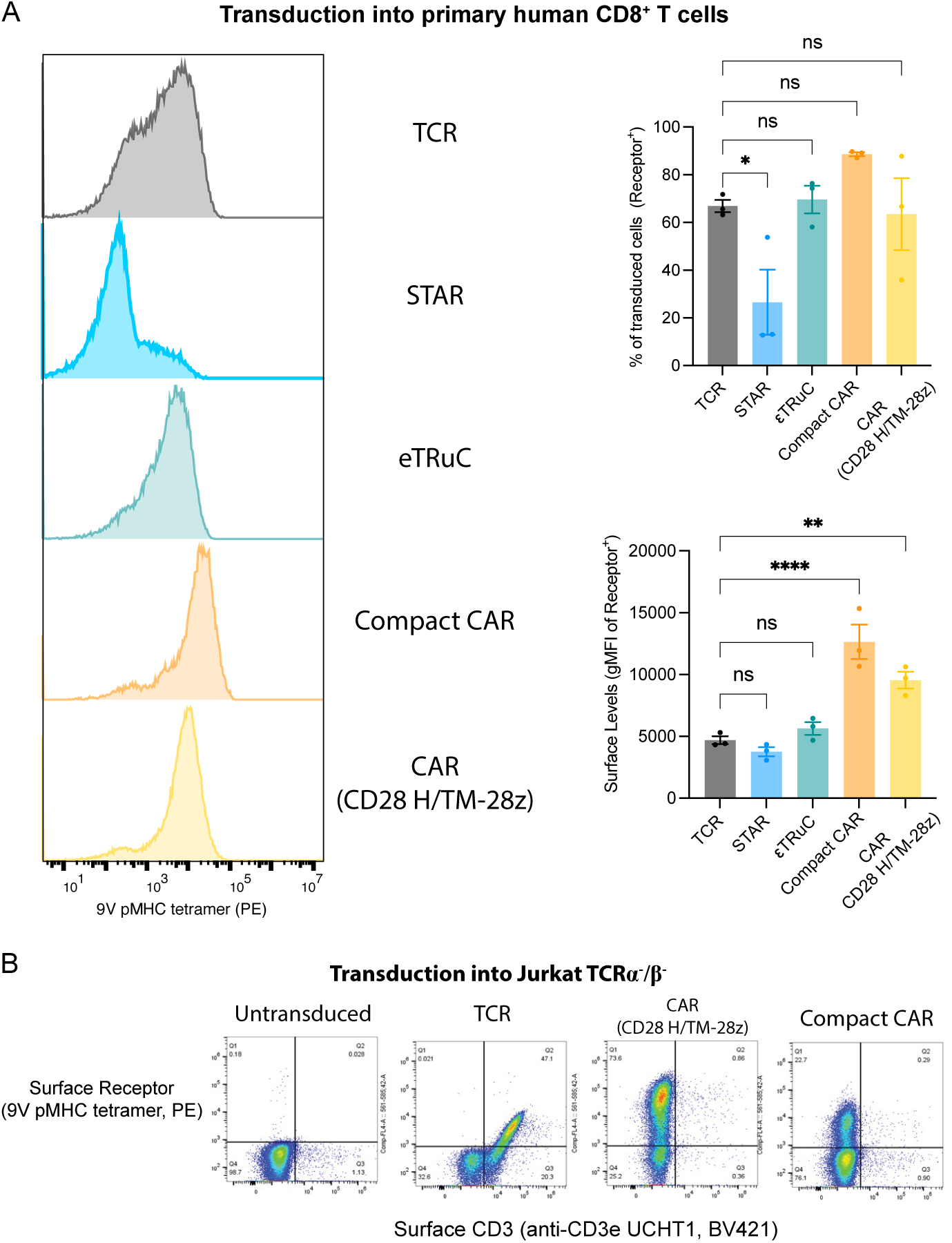
Surface expression of the Compact CAR is independent of the TCR-CD3 complex. **(A)** Quantitative analysis of surface expression of the indicated antigen receptors in primary human CD8+ T cells. The percent of T cells expressing the antigen receptor (transduction efficiency) was similar for all antigen receptors with the exception of the STAR (top right). The surface level of antigen receptors that associated with the CD3 complex were similar and lower than the Compact CAR and CAR (bottom right), which likely reflects competition for and/or limited components of the CD3 molecules. **(B)** Transduction of the TCR but not a standard CAR or the Compact CAR induces upregulation of surface CD3 on TCR*α*^−^*/β*^−^ Jurkat T cells. A t-test with Holm-Sidak multiple comparison correction is used to determine p-values. Abbreviations: * = p-value≤0.05, ** = p-value≤0.01, *** = p-value≤0.001, **** = p-value≤0.0001.

**Figure S3:**
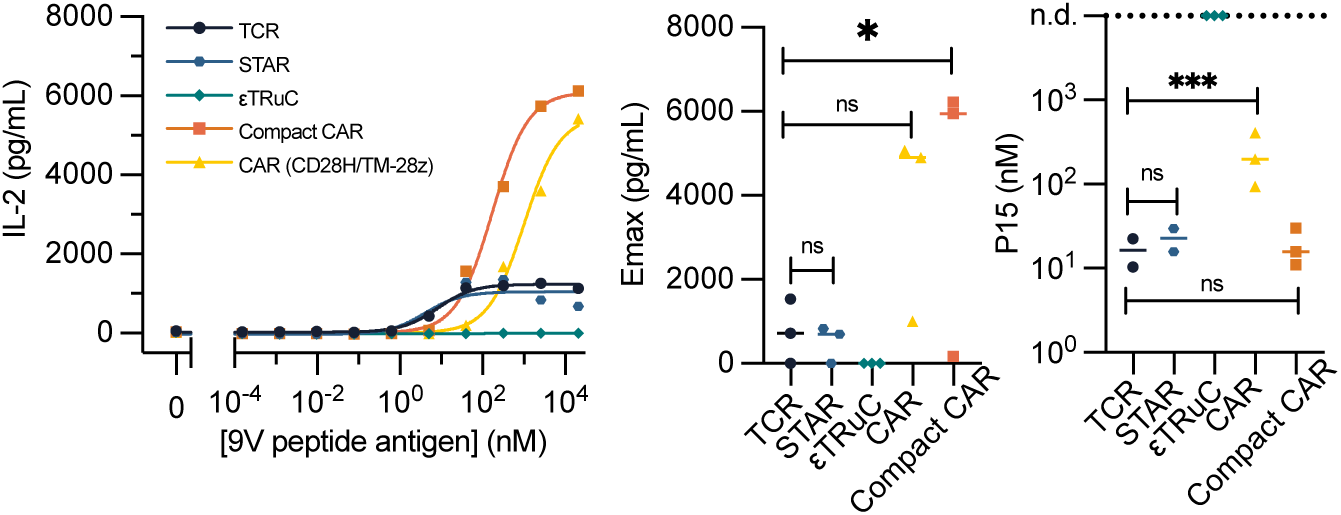
Additional measures of T cell activation for the experiment in. **Fig 1C-D**. Supernatant levels of IL-2. The eTRuC upregulated 4-1BB but did not produce IL-2 or IFNg. A t-test with Holm-Sidak multiple comparison correction is used to determine p-values. Abbreviations: * = p-value≤0.05, ** = p-value≤0.01, *** = p-value≤0.001, **** = p-value≤0.0001.

**Figure S4:**
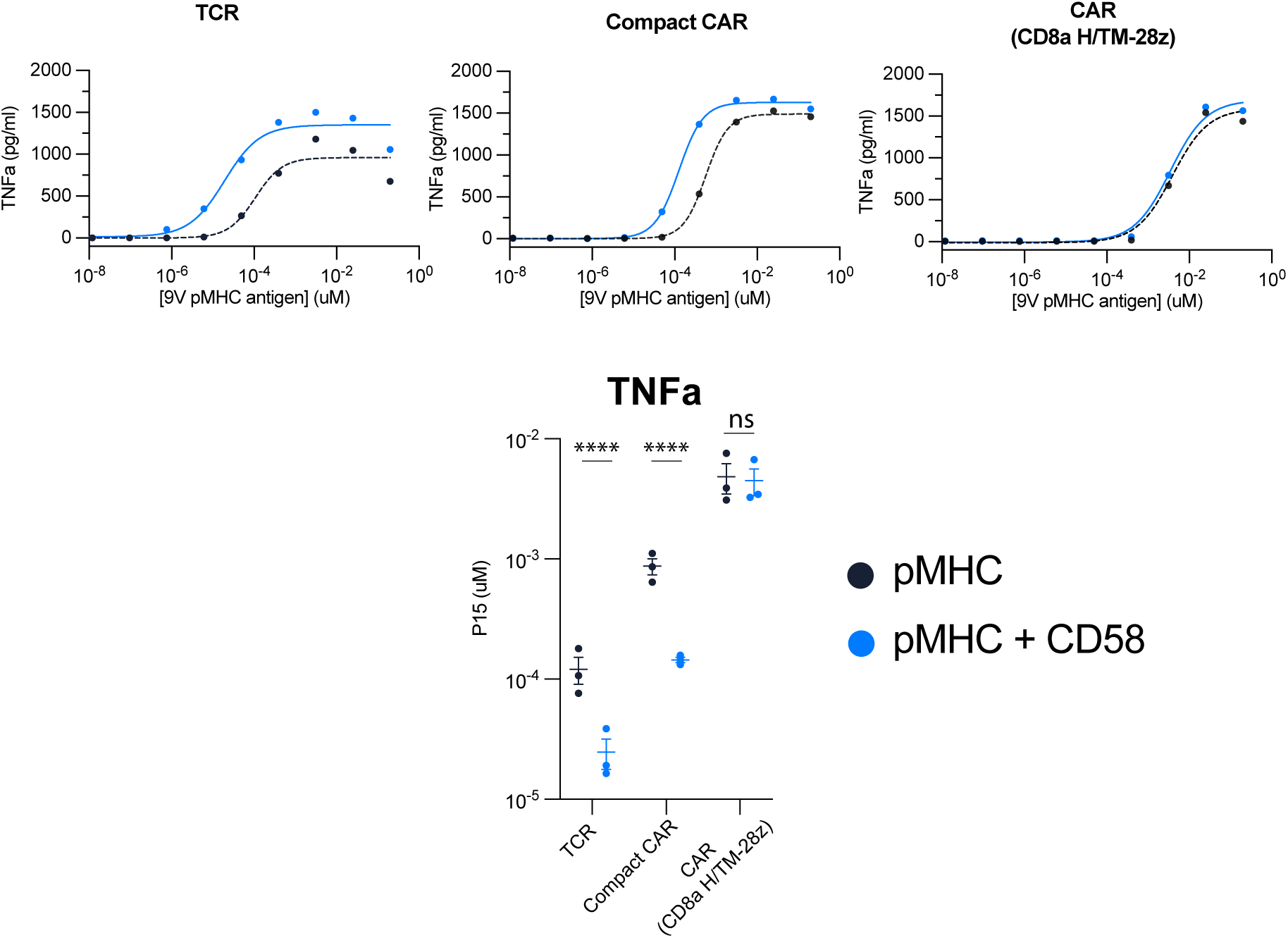
Additional measures of T cell activation for the experiment in. Fig 2. Representative dose-responses for supernatant levels of TNFa (top) with summary sensitivity measures across N=3 independent experiments (bottom). A t-test with Holm-Sidak multiple comparison correction is used to determine p-values. Abbreviations: * = p-value≤0.05, ** = p-value≤0.01, *** = p-value≤0.001, **** = p-value≤0.0001.

**Figure S5:**
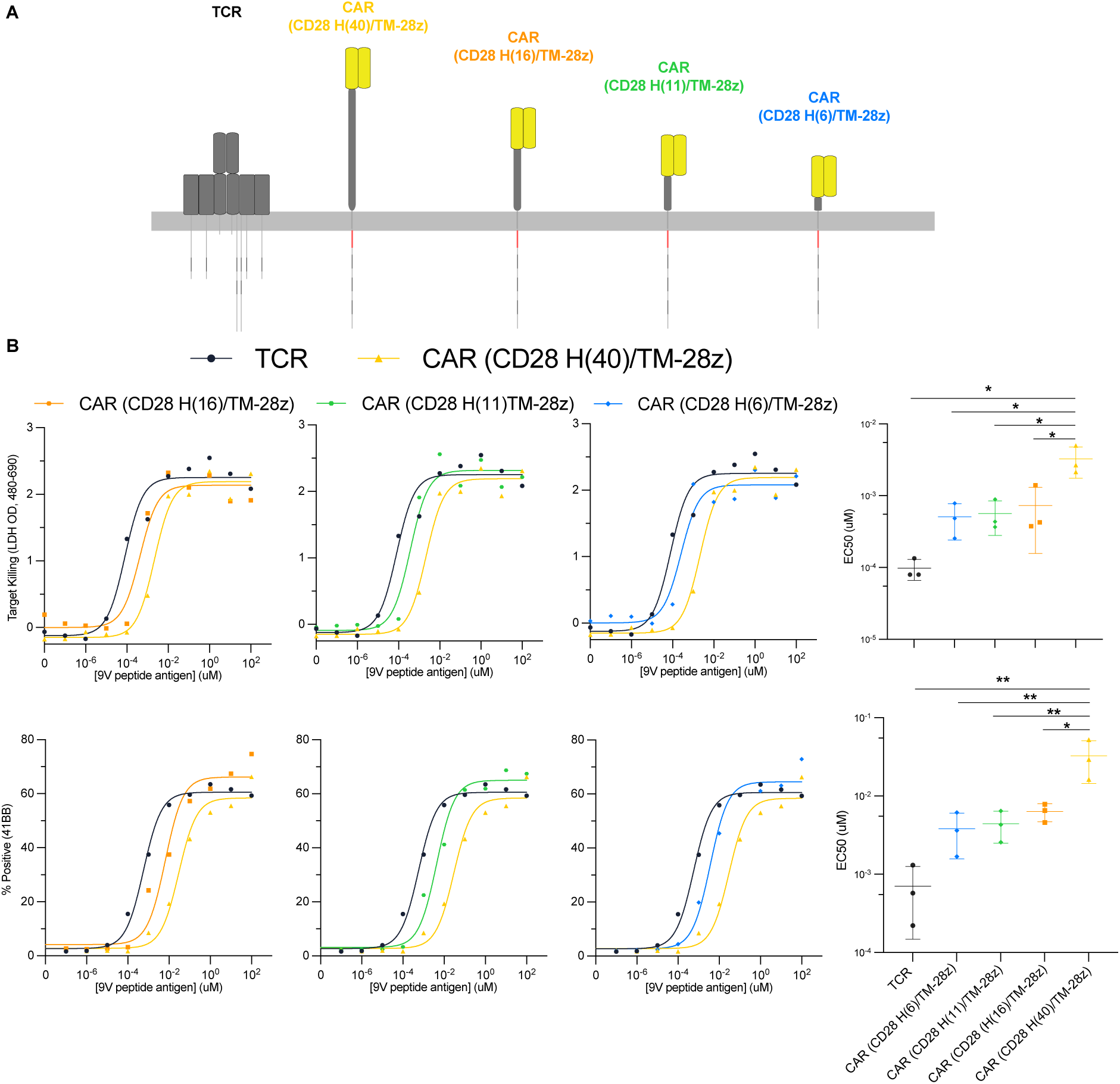
Reducing the length of the extracellular hinge of a CAR increases its antigen sensitivity. **(A)** Cartoon depicting the CD28 H/TM-z CAR containing a 40 amino acid hinge and truncated variants. **(B)** Representative dose-response and summary sensitivity measures across N=3 independent experiments for target cell killing (top) and the activation marker 4-1BB (bottom). The TCR and wild-type CAR are shown in each experiment for comparisons. A t-test with Holm-Sidak multiple comparison correction is used to determine p-values. Abbreviations: * = p-value≤0.05, ** = p-value≤0.01, *** = p-value≤0.001, **** = p-value≤0.0001.

**Figure S6:**
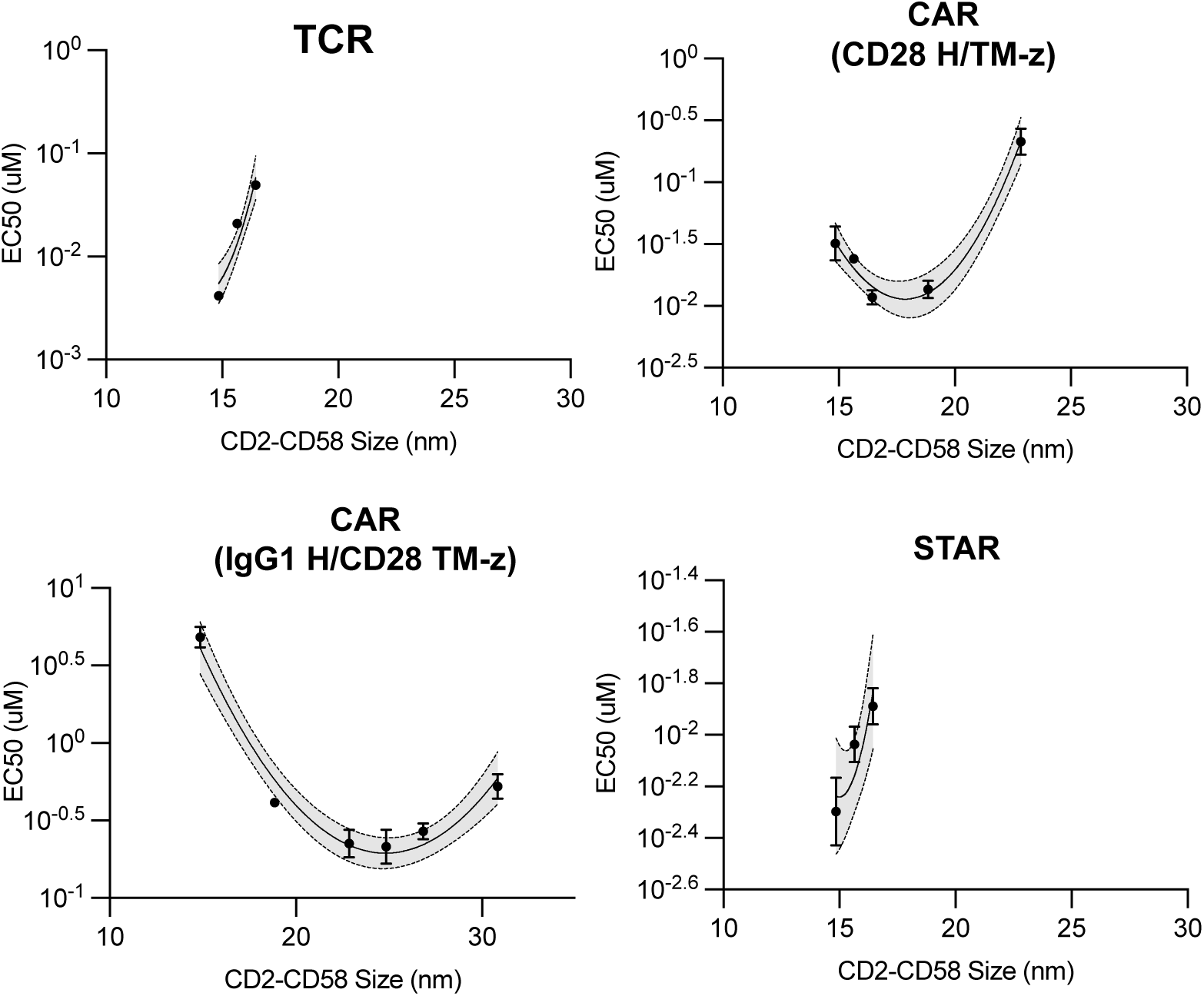
Estimating the preferred CD2-CD58 size for optimal antigen sensitivity. The antigen sensitivity (EC_50_) over the CD2-CD58 size (see Methods) is fitted to a parabolic function to determine the CD2-CD58 size that minimises EC_50_ (‘preferred CD2-CD58 size’).

**Figure S7:**
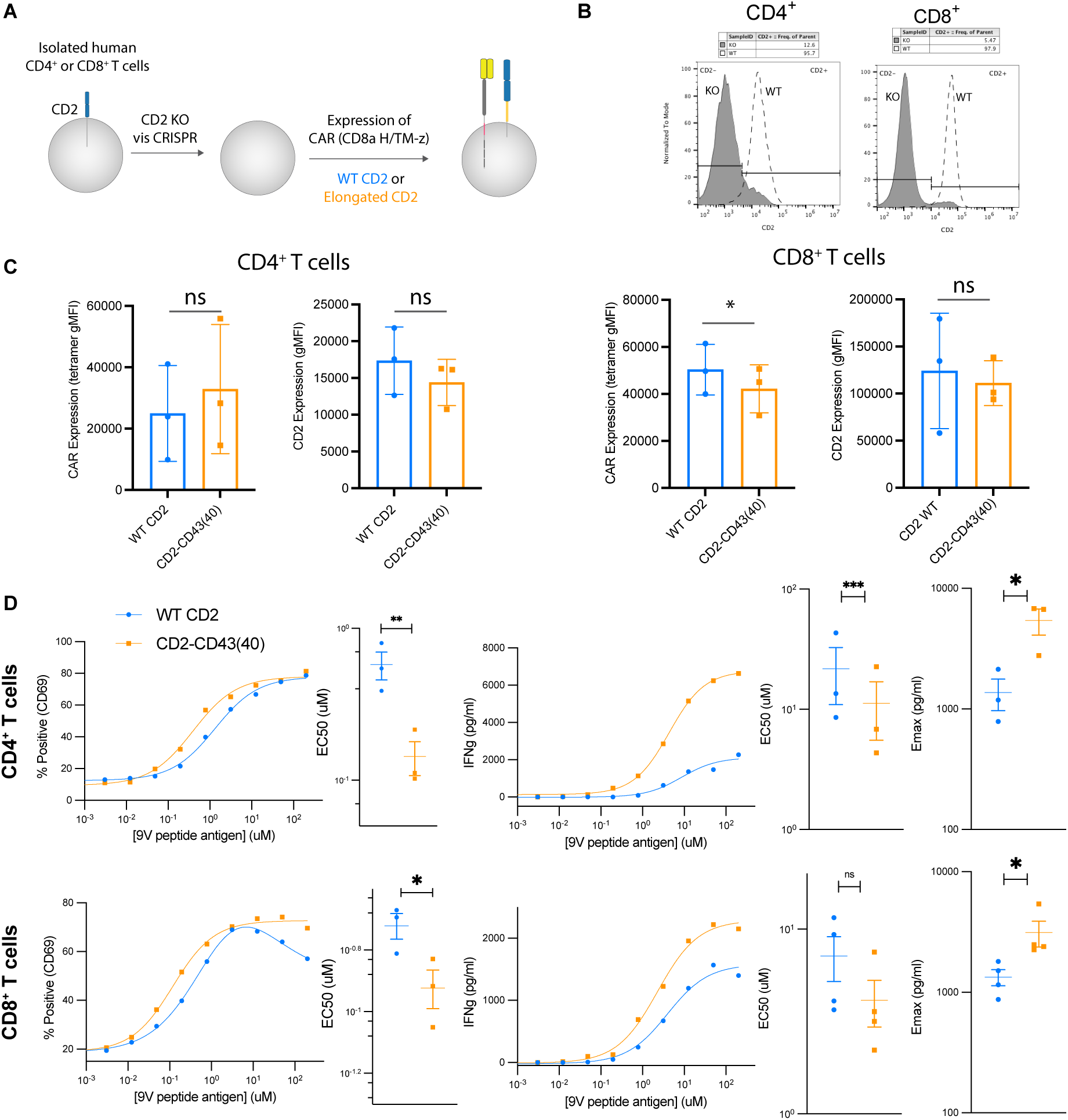
Primary human CD4+ and CD8+ CAR-T cells display enhanced sensitivity and efficacy with elongated CD2. **(A)** Primary T cells are genetically modified via CRISPR to knockout CD2 and then expanded and transduced with a plasmid encoding the CAR and either wild-type CD2 or elongated CD2. **(B)** Surface expression of CD2 before and after knockout of CD2 (before transduction of CAR and CD2). **(C)** Surface expression of the transduced CAR and CD2 on CD4+ T cells (left) and CD8+ T cells (right). **D** Representative dose-response and summary measures of T cell activation in response to U87 glioblastoma cells. A paired t-test is used to determine p-values across N=3 independent experiments. Abbreviations: * = p-value≤0.05, ** = p-value≤0.01, *** = p-value≤0.001, **** = p-value≤0.0001.

**Figure S8:**
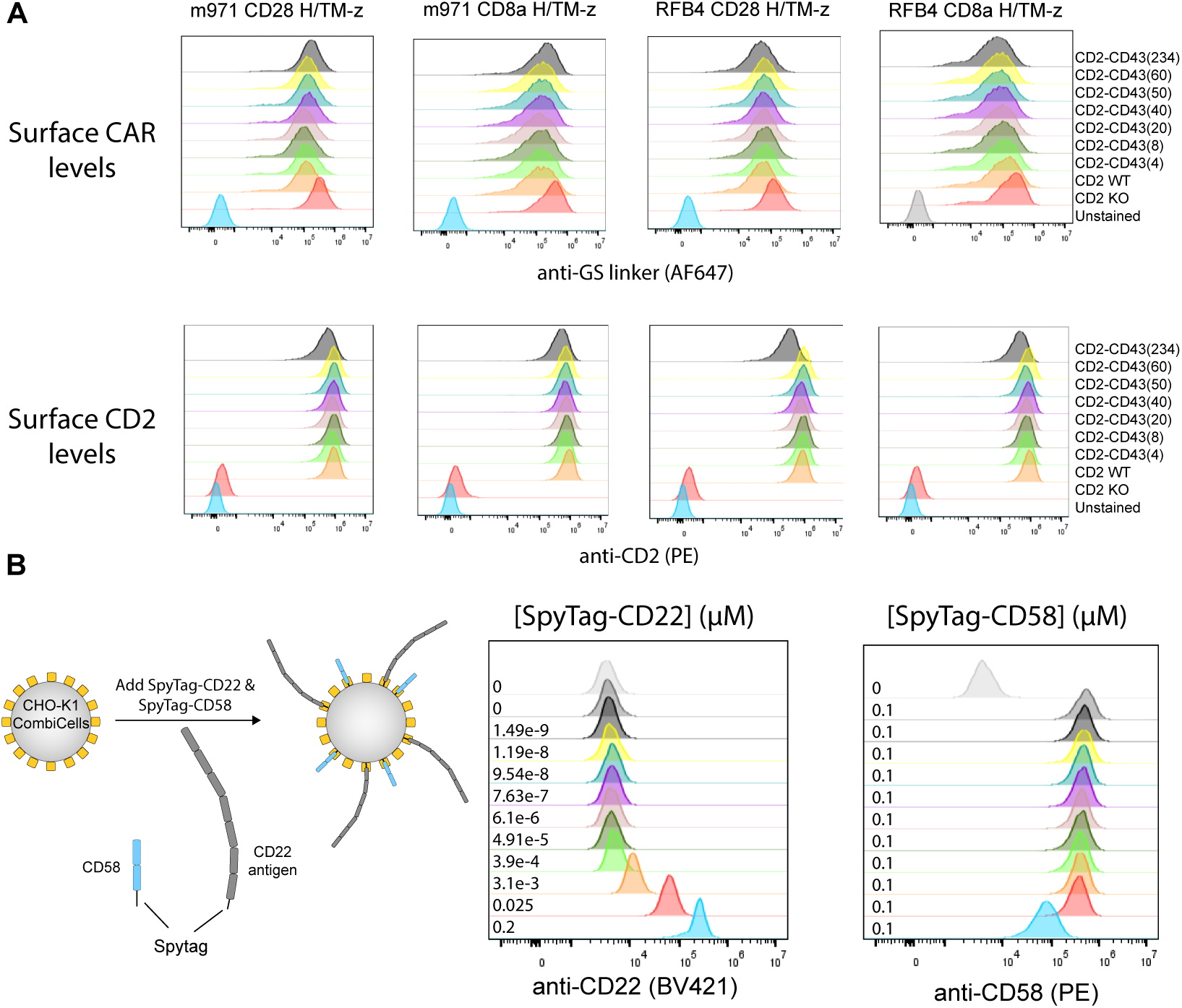
Surface expression of receptors and ligands (related to. Fig. 7**). (A)** Surface expression of CARs (top) and CD2 (bottom) display matched expression on the indicated Jurkat T cell line. **(B)** Surface expression of Spytag-CD22 and Spytag-CD58 on the surface of CHO-K1 CombiCells for the indicated concentration.

**Figure S9:**
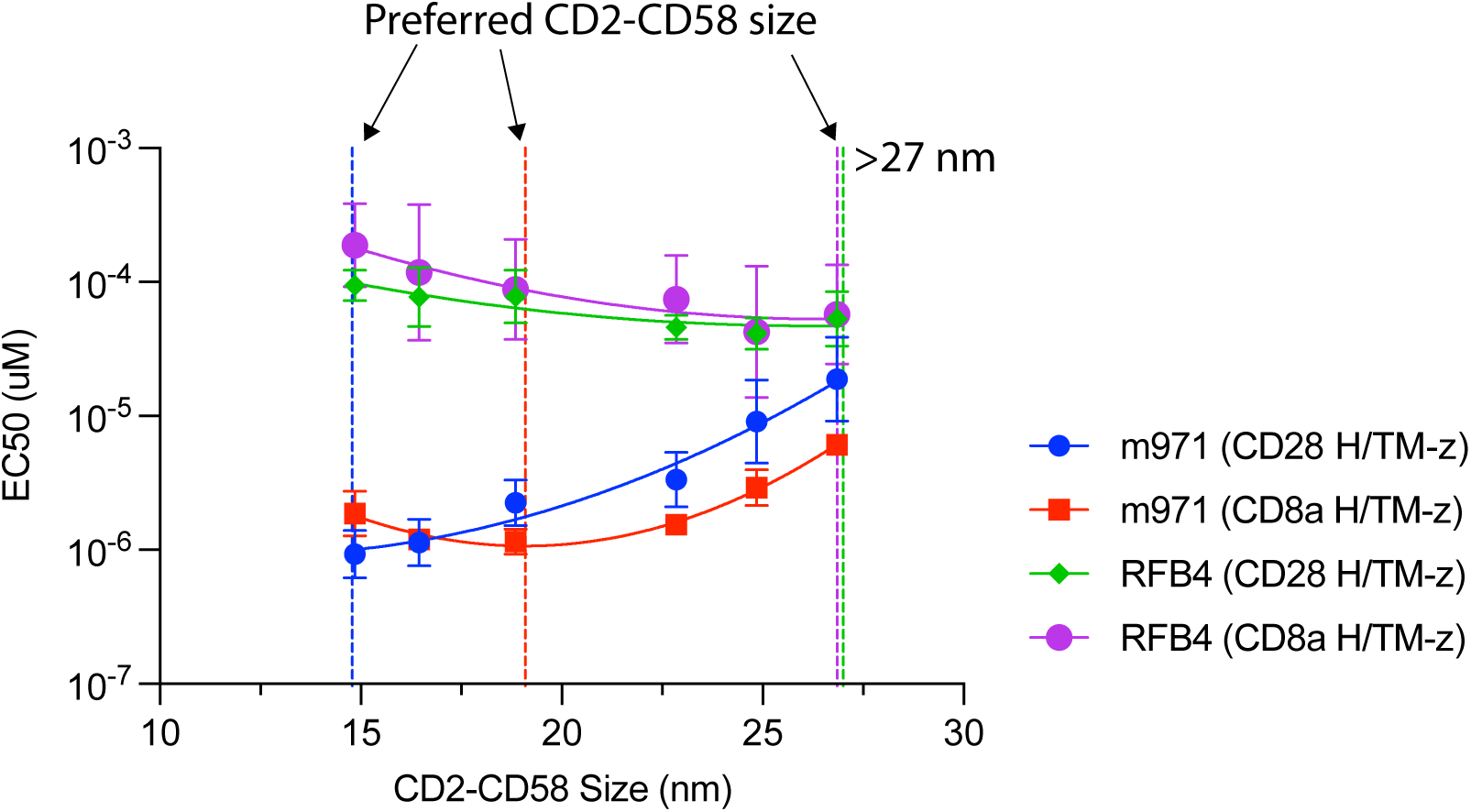
Estimating the preferred CD2-CD58 size for CD22 targeting CARs. The EC50 plotted over the estimated CD2-CD58 size. The preferred CD2-CD58 size is shown as a vertical dashed line for each antigen receptor. Data points represent mean±SEM for N=3 (m971 CD28/CD8a H/TM-z, RFB4 CD8a H/TM-z) and N=2 (RFB4 CD28 H/TM-z).

