## Supplementary Information for "Optimising CAR-T cell sensitivity by engineering matched extracellular sizes between CAR/antigen and CD2/CD58 adhesion complexes"

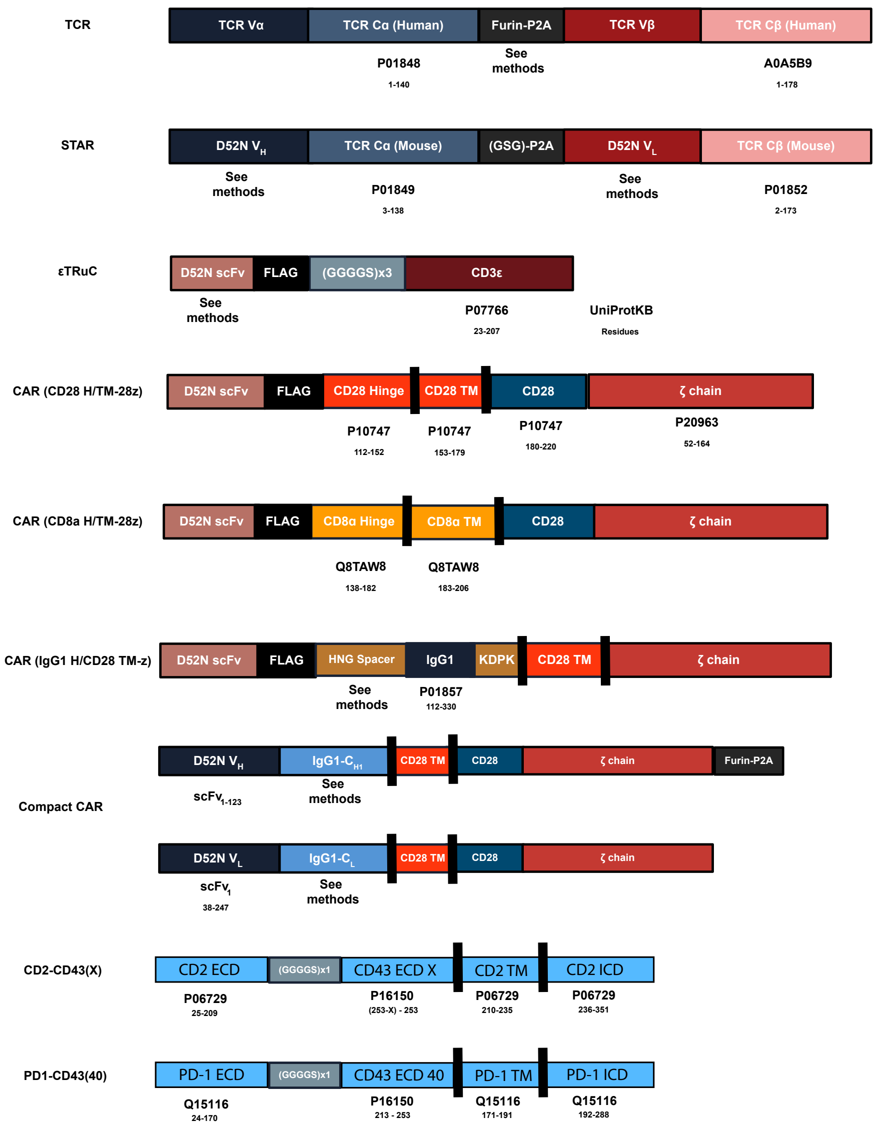


Figure S1: **Architecture and sequences of surface molecules used in the present study (also see Methods).**

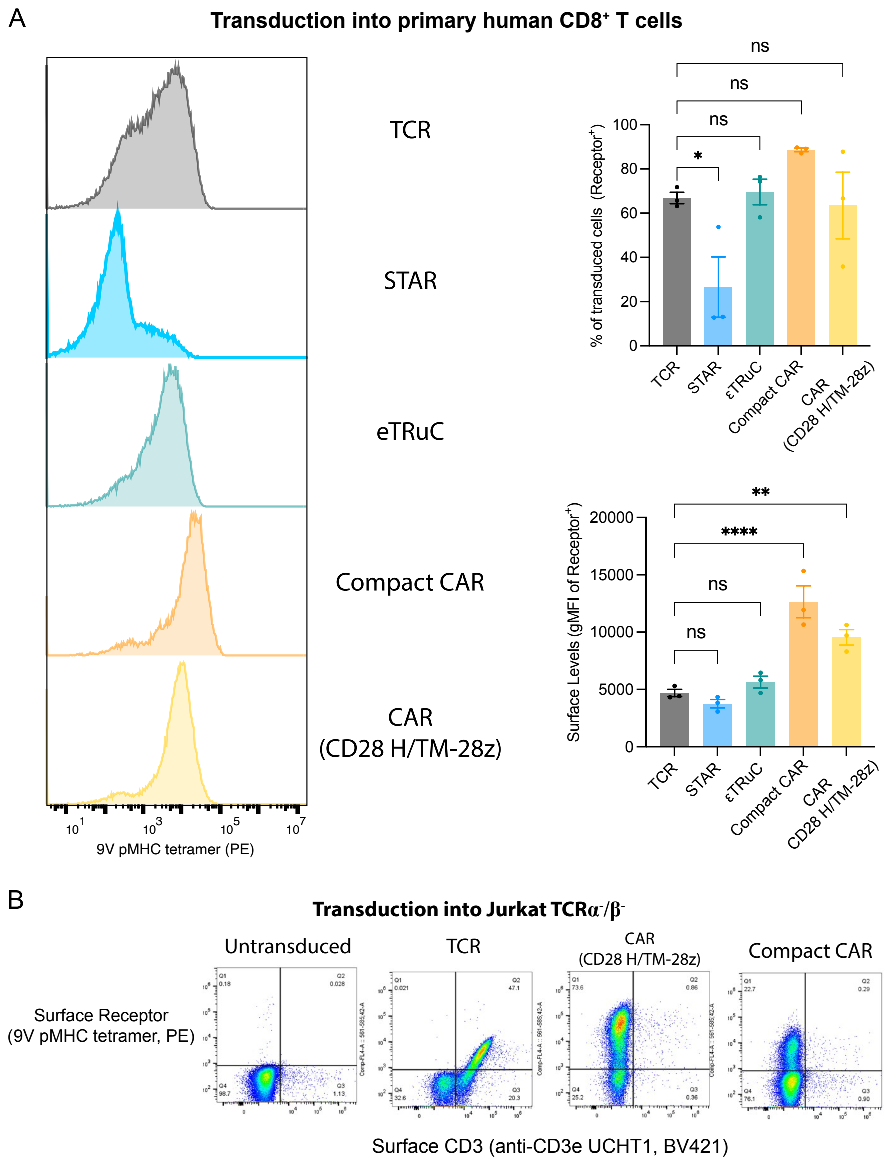


Figure S2: **Surface expression of the Compact CAR is independent of the TCR-CD3 complex.** **(A)** Quantitative analysis of surface expression of the indicated antigen receptors in primary human CD8+ T cells. The percent of T cells expressing the antigen receptor (transduction efficiency) was similar for all antigen receptors with the exception of the STAR (top right). The surface level of antigen receptors that associated with the CD3 complex were similar and lower than the Compact CAR and CAR (bottom right), which likely reflects competition for and/or limited components of the CD3 molecules. **(B)** Transduction of the TCR but not a standard CAR or the Compact CAR induces upregulation of surface CD3 on TCR$\alpha^-/\beta^-$ Jurkat T cells. A t-test with Holm-Sidak multiple comparison correction is used to determine p-values. Abbreviations: * = p-value$\leq$0.05, ** = p-value$\leq$0.01, *** = p-value$\leq$0.001, **** = p-value$\leq$0.0001.

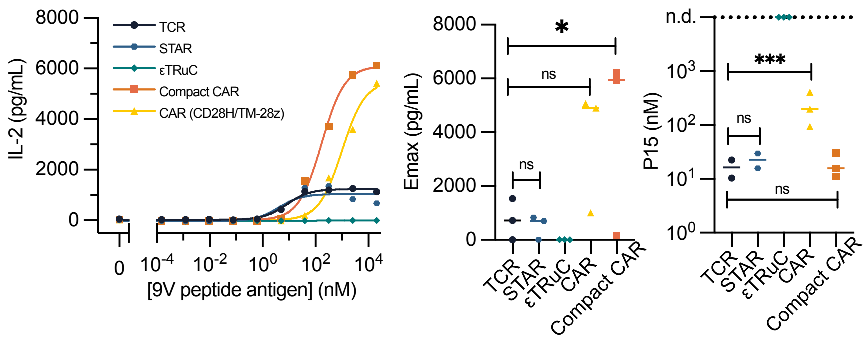


Figure S3: **Additional measures of T cell activation for the experiment in Fig 1C-D.** Supernatant levels of IL-2. The eTRuC upregulated 4-1BB but did not produce IL-2 or IFNg. A t-test with Holm-Sidak multiple comparison correction is used to determine p-values. Abbreviations: * = p-value≤0.05, ** = p-value≤0.01, *** = p-value≤0.001, **** = p-value≤0.0001.

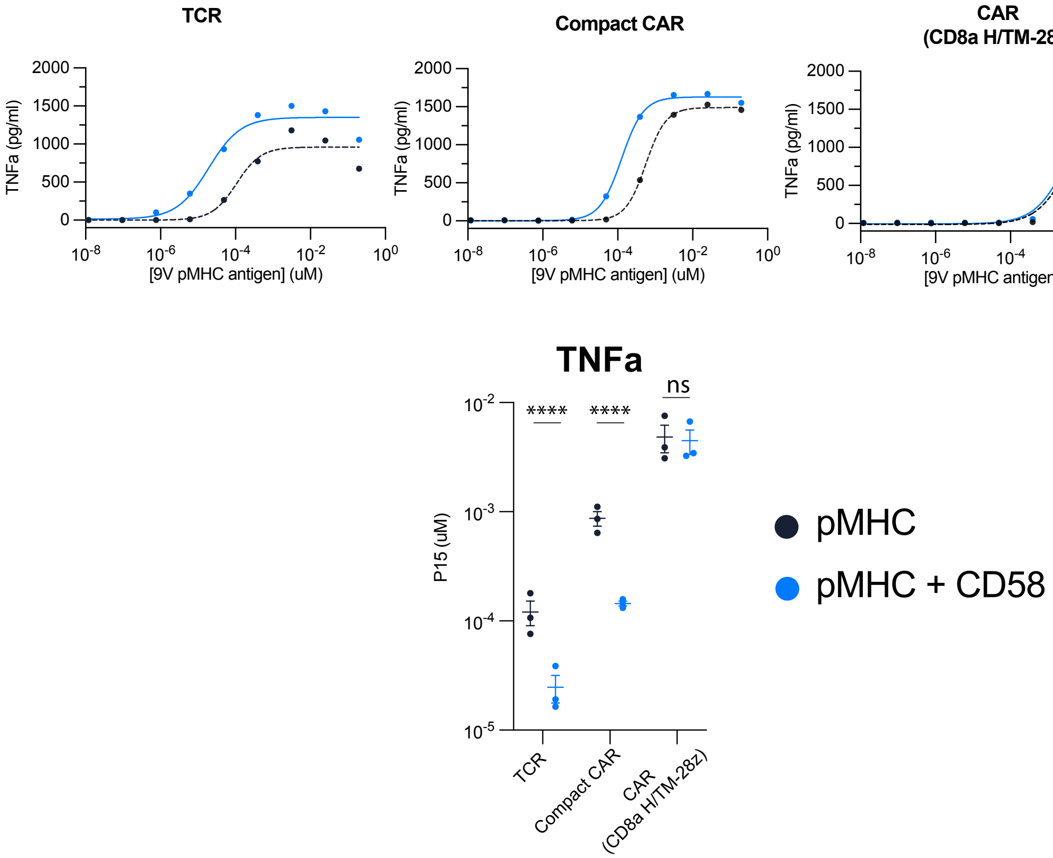


Figure S4: **Additional measures of T cell activation for the experiment in Fig 2.** Representative dose-responses for supernatant levels of TNFa (top) with summary sensitivity measures across N=3 independent experiments (bottom). A t-test with Holm-Sidak multiple comparison correction is used to determine p-values. Abbreviations: * = p-value≤0.05, ** = p-value≤0.01, *** = p-value≤0.001, **** = p-value≤0.0001.

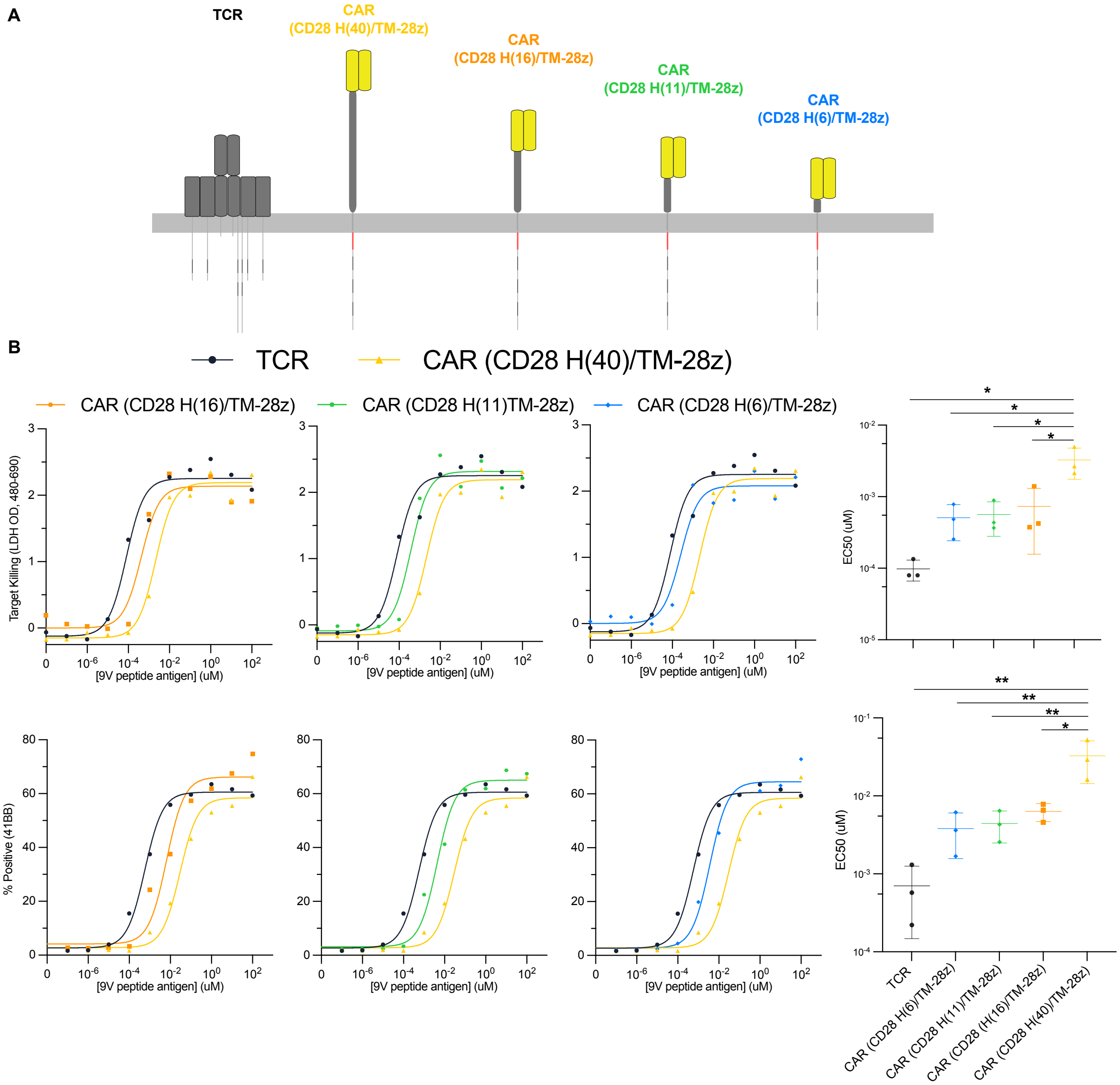


Figure S5: **Reducing the length of the extracellular hinge of a CAR increases its antigen sensitivity. (A)** Cartoon depicting the CD28 H/TM-z CAR containing a 40 amino acid hinge and truncated variants. **(B)** Representative dose-response and summary sensitivity measures across N=3 independent experiments for target cell killing (top) and the activation marker 4-1BB (bottom). The TCR and wild-type CAR are shown in each experiment for comparisons. A t-test with Holm-Sidak multiple comparison correction is used to determine p-values. Abbreviations: * = p-value$\leq$0.05, ** = p-value$\leq$0.01, *** = p-value$\leq$0.001, **** = p-value$\leq$0.0001.

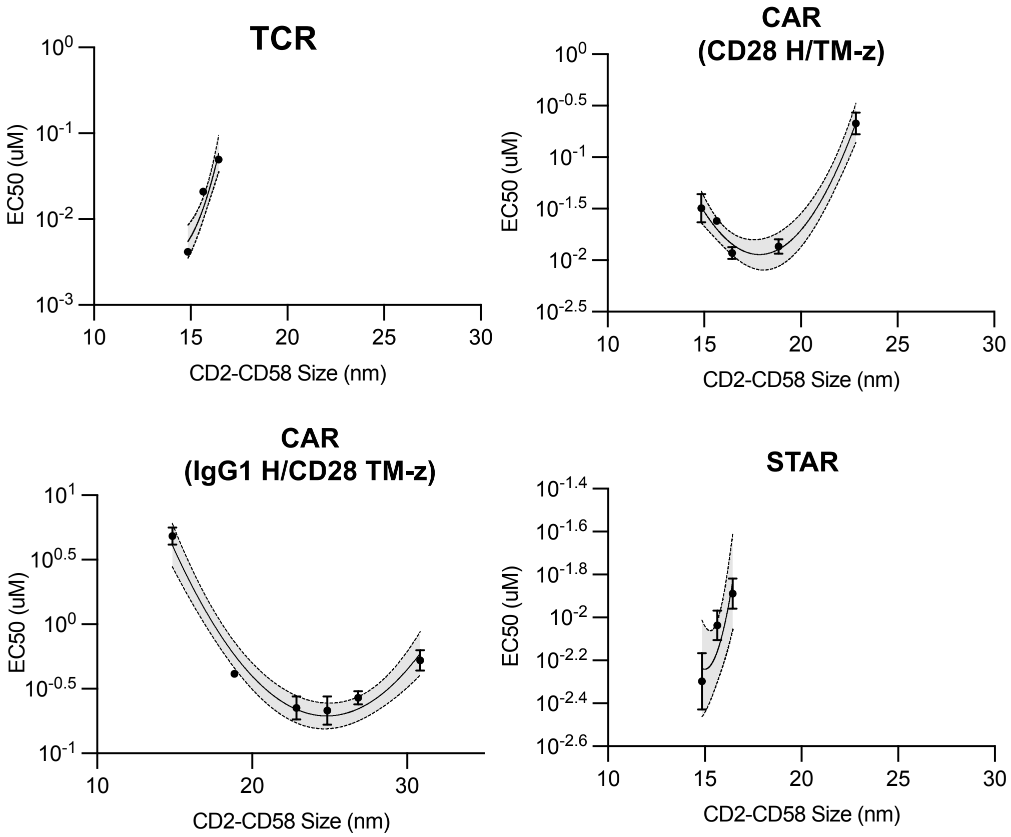


Figure S6: **Estimating the preferred CD2-CD58 size for optimal antigen sensitivity.** The antigen sensitivity ($EC_{50}$) over the CD2-CD58 size (see Methods) is fitted to a parabolic function to determine the CD2-CD58 size that minimises $EC_{50}$ ('preferred CD2-CD58 size').

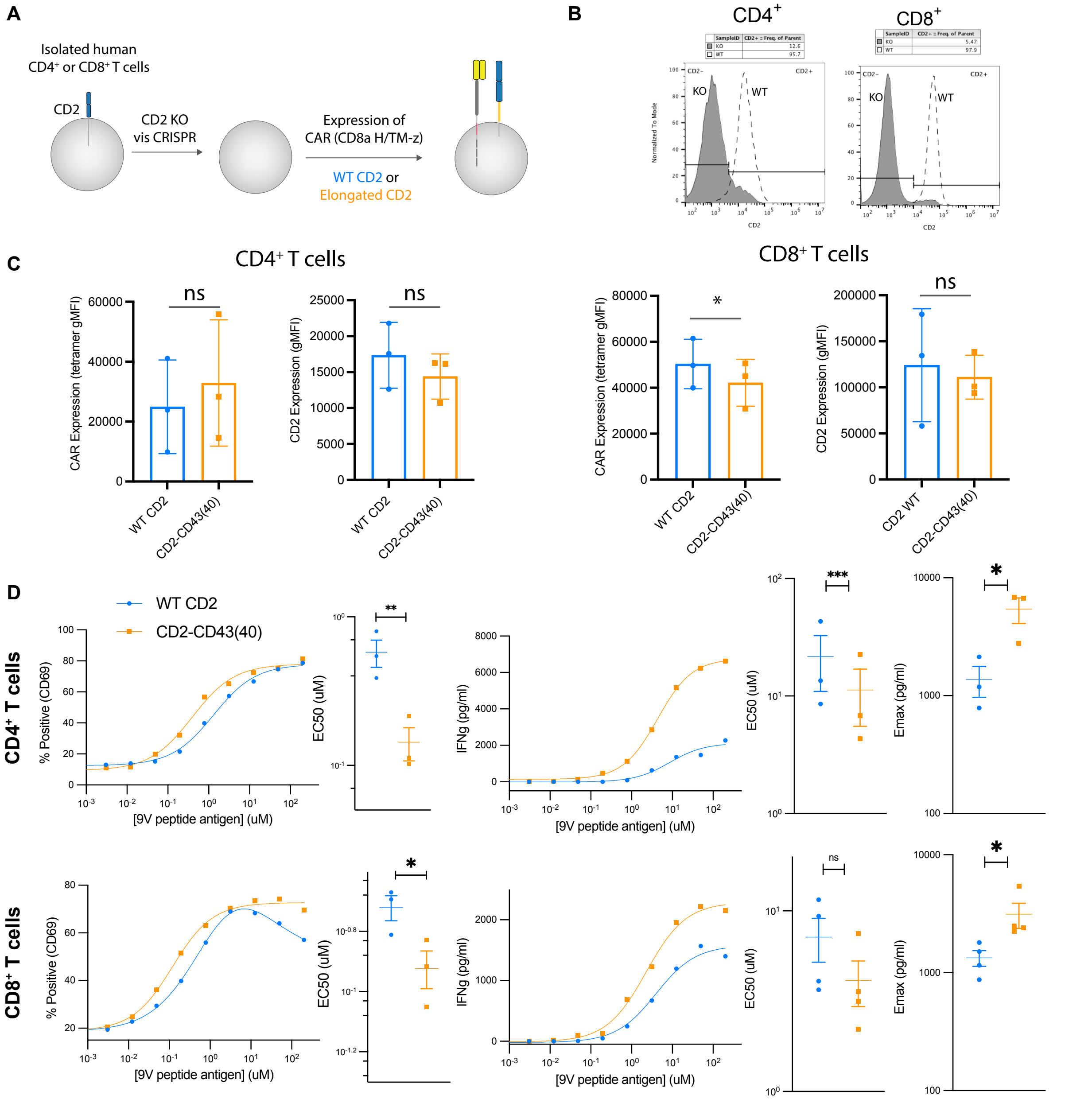


Figure S7: **Primary human CD4+ and CD8+ CAR-T cells display enhanced sensitivity and efficacy with elongated CD2.** **(A)** Primary T cells are genetically modified via CRISPR to knockout CD2 and then expanded and transduced with a plasmid encoding the CAR and either wild-type CD2 or elongated CD2. **(B)** Surface expression of CD2 before and after knockout of CD2 (before transduction of CAR and CD2). **(C)** Surface expression of the transduced CAR and CD2 on CD4+ T cells (left) and CD8+ T cells (right). **D** Representative dose-response and summary measures of T cell activation in response to U87 glioblastoma cells. A paired t-test is used to determine p-values across N=3 independent experiments. Abbreviations: * = p-value≤0.05, ** = p-value≤0.01, *** = p-value≤0.001, **** = p-value≤0.0001.

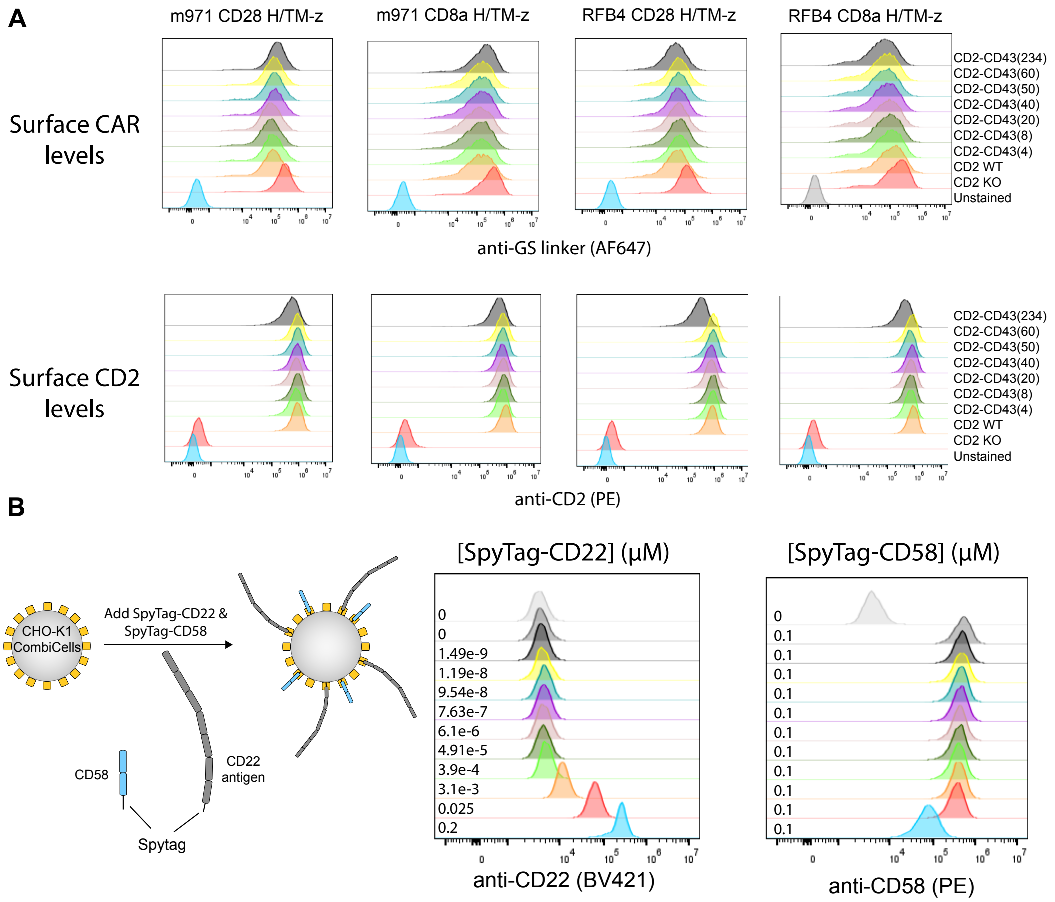


Figure S8: **Surface expression of receptors and ligands (related to Fig. 7).** **(A)** Surface expression of CARs (top) and CD2 (bottom) display matched expression on the indicated Jurkat T cell line. **(B)** Surface expression of Spytag-CD22 and Spytag-CD58 on the surface of CHO-K1 CombiCells for the indicated concentration.

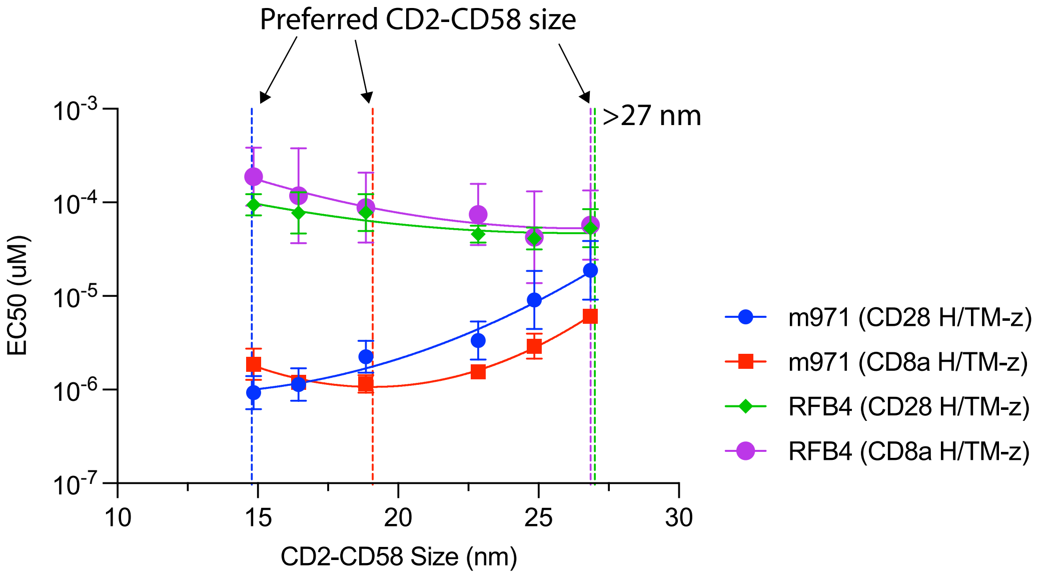


Figure S9: **Estimating the preferred CD2-CD58 size for CD22 targeting CARs.** The EC50 plotted over the estimated CD2-CD58 size. The preferred CD2-CD58 size is shown as a vertical dashed line for each antigen receptor. Data points represent mean±SEM for N=3 (m971 CD28/CD8a H/TM-z, RFB4 CD8a H/TM-z) and N=2 (RFB4 CD28 H/TM-z).
